# LRRK2-G2019S Synergizes with Ageing and Low-Grade Inflammation to Promote Gut and Peripheral Immune Cell Activation that Precede Nigrostriatal Degeneration

**DOI:** 10.1101/2022.09.01.505977

**Authors:** Carmela Giachino, Cataldo Tirolo, Salvatore Caniglia, Maria F. Serapide, Francesca L’Episcopo, Federico Bertoli, Claudio Giuliano, Marika Mearelli, Meike Jakobi, Nicole Schneiderhan-Marra, Michela Deleidi, Bianca Marchetti

**Affiliations:** OASI-Research Institute-IRCCS, Neuropharmacology Laboratory, Troina-EN-Italy; Biomedical and Biotechnological Sciences, Pharmacology Section, University of Catania-Italy; Mitochondria and Inflammation in Neurodegenerative Diseases, DZNE, Tübingen-Germany; Hertie Institute for Clinical Brain Research, University of Tübingen; NMI Natural and Medical Sciences Institute, University of Tübingen, Reutlingen, Germany

**Keywords:** LRRK2, low-grade inflammation, ageing, Parkinson’s disease, nigrostriatal dopaminergic degeneration, colonic α-synuclein, central and peripheral inflammation, brain-gut-axis

## Abstract

**Background:** Mutations in the leucine-rich repeat kinase 2 (*LRRK2*) gene are the most frequent cause of familial Parkinson’s disease (PD). The incomplete penetrance of *LRRK2* mutations suggest that additional hits are required for disease onset. We hypothesized that chronic low-grade inflammation interacts with LRRK2 G2019S, the most frequent PD-associated mutation, to activate peripheral and central immune reactions and drive age-dependent neurodegeneration.

**Methods and Results:** We exposed wild-type and LRRK2 G2019S mice to a low chronic dose of lipopolysaccharide, and we performed a longitudinal analysis of central and peripheral immune reactions and neurodegeneration. Low-dose inflammation triggered nigrostriatal degeneration, macrophage/monocyte brain infiltration, and astro-/microgliosis. LRRK2 G2019S mice showed an early dysregulation of peripheral cytokines, increased CD4^+^ T-cell infiltration and α-synuclein aggregation in the colon. Interestingly, peripheral immune activation and colonic α-synuclein aggregation precede astro-/microgliosis and neurodegeneration.

**Conclusions:** Our study suggests an early role of the peripheral immune system and the gut in LRRK2 PD and provides a novel model to study early therapeutic immune targets and biomarkers.

## 1 BACKGROUND

Parkinson’s disease (PD) is the most prevalent neurodegenerative movement disorder, mostly occurring in a sporadic manner, through a complex interaction between genetic risk, ageing, and environmental exposure.^1,2^ Progressive loss of midbrain dopaminergic neurons (mDAns) in the substantia nigra pars compacta (SNpc) along with the presence of Lewy bodies and Lewy neurites composed of α-synuclein (α-syn) in the SNpc and other brain and peripheral tissues are the main neuropathological hallmarks of the disease.^3–7^ Degeneration of mDAn terminals in the striatum (Str) is slow in most cases, but irreversible, with current therapies providing only symptomatic relief.^8^

Among genetic risk factors, mutations in the leucine-rich repeat kinase 2 (*LRRK2*) gene are the most prevalent cause of late-onset autosomal dominant PD.^9,10^ Both idiopathic and LRRK2 PD display a strong age-dependent development of motor and non-motor symptoms and are clinically indistinguishable.^11,12^ The incomplete penetrance of *LRRK2* mutations is recognized in humans and LRRK2 murine models.^12–14^ Notably, transgenic mice overexpressing pathogenic *LRRK2* mutations fail to develop an overt neurodegenerative phenotype—up to 2 years of age, which has hampered the identification of key mechanisms that drive neurodegeneration.^15–21^ These findings suggest that the disease may result from a complex interplay between genetic predisposition and exogenous insults, such as the ageing process, inflammation, and toxic exposure. ^22–26^

In fact, the degeneration of mDAns in the SNpc also occurs in healthy ageing,^23,27–30^ and *LRRK2* penetrance progressively increases with age.^13,14,31,32^ Both ageing and inflammation are recognized as key factors contributing to both sporadic and familial PD.^26,33–37^ Furthermore, burgeoning evidence shows that PD involves the gut before affecting the brain, leading to the fascinating hypothesis that an age-dependent dysfunction of the gut-brain axis might be involved in disease initiation.^38–41^ Changes in the crosstalk between the central and the peripheral immune system^14,41,42^ may exacerbate glial reactions and promote nigrostriatal neuron vulnerability/death. In this regard, reactive astrocytes, microglia, and infiltrated peripheral immune cells play major roles in immune-related mDAn degeneration.^33–35,42–49^ LRRK2 is highly expressed in both innate and adaptive immune cells, being tightly regulated by inflammatory triggers.^37,50–52^ Interestingly, *LRRK2* is an IFN-γ target gene and it is involved in the host immune response to pathogens.^50,51,53–56^ Significantly, LRRK2 has also been linked to inflammatory bowel diseases and several bacterial infections, including enteric pathogens.^57–61^ Hence, LRRK2 may be involved in the regulation of the peripheral immune system, including intestinal immunity, and, potentially, colonic α-syn pathology. Whether LRRK2-related intestinal and peripheral immune processes precede neurodegeneration is still unknown.

We herein hypothesized that a chronic low-grade systemic inflammatory stimulus interacts with *LRRK2* G2019S (the most frequent PD-associated LRRK2 mutation) as the mice age to alter the crosstalk between the peripheral and central immune system and to promote nigrostriatal degeneration. To address this hypothesis, we developed a mouse model integrating LRRK2 genetic vulnerability, low-grade inflammation, and ageing, the key risk factors for PD.^2,22,23,26,33,47,61–64^

It is well known that aged animals show higher pro-inflammatory cytokines levels, both in the brain and in the periphery, in response to systemic acute immune activation, when compared to younger animals.^65–69^ As ageing is recognized as the primary risk factor influencing the clinical presentations and the progression of both sporadic and LRRK2 PD, ^2,22,23,26^ we assessed whether *LRRK2* G2019S mutation combined with chronic systemic moderate inflammation interacts with the ageing process to modulate systemic and central immune reactions as well as nigrostriatal neurodegeneration. The ageing process is relevant as a single acute injection of a high dose (5 mgkg^−1^) of lipopolysaccharide (LPS), which is commonly used to study the effect of endotoxemia, results in a moderate (÷-25-30%) loss of mDAns between 7 days and 7 months (M) post-injection in transgenic *LRRK2* R1441G mutant mice.^70^ Furthermore, in this experimental setup, neuroinflammatory processes observed in *LRRK2* mutant mice are mostly mediated by circulating inflammatory molecules.^70^ Similarly, the study of Litteljohn and co. (2018)^71^ shows that acute systemic LPS injection is not sufficient to induce microgliosis in young *LRRK2* G2019S mice, suggesting that other factors, such as ageing, may interact with the *LRRK2* genotype in the regulation of the inflammatory processes.

To mimic a chronic, low-grade inflammation, we exposed both young and middle-aged WT and G2019S mice to a very low LPS dose (0.1 mg kg^−1^), administrated twice weekly for 12 weeks. We selected two previously well-characterized temporal windows-young-adult and middle-aged, ^67–69,73–76^ as during middle-age nigrostriatal DA neuron vulnerability progressively increases, while dopaminergic adaptive/neurorepair capacity diminishes.^28–30,72–75^ To monitor the interaction between LRRK2 G2019S, low-grade inflammation and ageing, the mice were sacrificed at 6 M, 10 M, and 16 M.

We herein show that *LRRK2* G2019S synergizes with ageing and low-grade inflammation to trigger nigrostriatal DA degeneration. Specifically, *LRRK2* G2019S, low-grade inflammation, and ageing promote macrophage/monocyte infiltration in the CNS, CD4^+^ T-cell infiltration, and α-syn aggregation in the colon that precede mDAn degeneration. This new preclinical model will enable the identification of novel immune-related targets involved in mDAn degeneration, both at the central and peripheral level, and it will be useful to explore the early pathological events with clinical relevance and the development of disease-modifying therapies.

## 2 METHODS

The detailed methods for cell culture conditions, Western blot analysis, enzyme-linked immunosorbent assay (ELISA), quantitative-polymerase chain reaction (q-PCR), list of probes, and primary antibodies are available in the *Supplemental Materials*.

### 2.1 ANIMALS

Males C57BL/6J wild type (WT) and C57BL/6J LRRK2*G2019S 2AMjff/J transgenic (TG-G2019S) (stock 018785) mice purchased from Jackson Laboratory (Bar Harbor, ME, USA) at 8-12 weeks of age were housed at the animal facilities unit (C.A.Pi.R. Via Santa Sofia 97, Catania 13/2017-UT) and maintained on a 12:12-h light/dark cycle with ad libitum food and water. The animals were given 3 weeks to acclimate to the housing conditions prior to study commencement. The research project (authorization n. 345/2019-PR released by the Italian Ministry of Health 06-05-2019) and all experimental procedures were approved by the University Institutional Animal Experimentation and Ethical Committee (OPBA, University of Catania) and performed in accordance with NIH Guide for the Care and Use of Laboratory Animals. Experiments were carried out in accordance with The Code of Ethics of the World Medical Association (Declaration of Helsinki) for animal experiments.

### 2.2 EXPERIMENTAL DESIGN

WT and TG-G2019S mice were exposed to intraperitoneal (ip) injections of a low dose of lipopolysaccharide LPS (0,1 mg kg^−1^ Escherichia coli serotype O111:B4) (Sigma-Aldrich), administrated twice a week for 12 weeks (*Supplementary Figure 1*). Treatments were performed in two different age groups, 3M (young adult) and 7M (at the start of middle-age). The dose of LPS was selected based upon previous research, including our own, demonstrating only a transient, mild inflammatory response in young adult (3M-old) as opposed to the exaggerated inflammatory response observed in aged (≥ 20 M-old) mice.^65–68,76^ WT and TG mice exposed to ip injections of 0.9% sterile NaCl were used as controls. Mice (n = 20/experimental group) were randomly assigned to one of seven experimental conditions for each genotype: **1**. 3 M Basal (no injections, T=0); **2**. 6 M NaCl (NaCl injections started at 3 M); **3**. 6 M-LPS (LPS injections started at 3 M); **4**.10 M-NaCl (NaCl injections started at 7 M); **5**.10 M-LPS (injections started at 7M); **6**. 16 M NaCl (injections started at 7 M); **7**. 16 M LPS (injections started at 7 M). Clinical evaluation (body weight, mantel status, lethargy, reluctance to move, grooming behavior) was carried out weekly until sacrifice.

### 2.3 MOTOR BEHAVIOR ANALYSIS WITH THE ROTAROD

To test motor performance, we used the accelerated Rotarod (five-lane accelerating rotarod; Ugo Basile, Comerio, Italy), a motor behavioral test widely used to assess motor deficit in neurodegenerative disease models in rodents.^77^ Mice have to keep their balance on a horizontal rotating rod (diameter, 3 cm) and rotation speed is progressively increased every 30 sec by 4 rpm.^72^ To start the trials, the mice (five mice are tested at the same time) are placed on a rotating rod; when the mice fall down or when 5 min are completed, a switch is activated that automatically stops a timer. On the day of testing, the mice perform 5 trials, separated by an interval of 30 min between each trial. Before the test, WT and G2019S within the different treatment groups were housed five per cage and acclimated to a 12h shift in light/dark cycle so that the exercise occurred during the animal normal wake period. The Rotarod performance was assessed at day −7, 0 days, and during NaCL/LPS injection paradigm, at monthly intervals, for the determination of motor deficit (*Supplementary Figure 1*).

### 2.4 TISSUE HANDLING

For immunohistochemistry, mice (5-6 mice/genotype/age-group/treatment) were anesthetized and transcardially perfused with 0.9% NaCl, followed by 4 % paraformaldehyde in phosphate buffer (pH 7.2 at 4°C), the brains and peripheral tissues carefully removed and processed for immunofluorescent staining as described. Blood was collected into EDTA-treated tubes and serum was separated by centrifugation and stored at −20C. For gene expression and protein analysis, animals were sacrificed by cervical dislocation, the brains and peripheral tissues were quickly isolated and then stored at −80°C until assayed.

### 2.5 IMMUNOHISTOCHEMISTRY AND IMAGE ANALYSIS

For brain tissue analysis, serial coronal sections (14 μm-thick), encompassing the striatum (Bregma 1.54 to bregma −0.46) and the SNpc (Bregma −2.92 to bregma −3.8 mm) according to Franklin and Paxinos (1997)^78^ were collected, mounted on poly-L-lysine-coated slides, and processed as previously described.^73–75^ TH immunoreactivity was also detected using biotinylated secondary antibodies (Vector Laboratories) and diaminobenzidine (DAB, Vector Laboratories) as the developing agent. Cresyl violet was used to visualize Nissl substance. DA neuronal counts were performed by serial section analysis of the total number of TH^+^ neurons in the right and left SNpc through the entire extent of the SNpc using DAPI or PI as nuclear markers, as previously described.^72^ TH^+^ cells were counted through the entire rostro-caudal axis of the murine SNpc (bregma coordinates: 2.92, 3.08, 3.16, 3.20, 3.40, and 3.52) according to Franklin and Paxinos^78^ (1997) as described.^74,79^ Striatal TH- and dopamine transporter (DAT)-immunofluorescent (IF) fiber staining was assessed in n = 3 coronal sections at three levels (bregma coordinates: + 0.5, + 0.86, and 1.1 mm, respectively) of caudate-putamen (CPu), in n 5-6 mice/group/time. Cell counts were obtained for IBA1^+^/Dapi^+^ or Mac-1^+^/Dapi^+^ reactive microglial cells and GFAP^+^/Dapi^+^ astrocytes, averaged for each animal and the mean number of cells per mm^2^ per animal was estimated. A comparable countable area ranging from 1.90 mm^2^ to 2.00 mm^3^ was analyzed in the different groups. Results are expressed as % of saline-injected controls.

For intestine analysis, the tissue was fixed in 1% paraformaldehyde for 2 hours, washed with 50 mmol/L NH4Cl, and cryoprotected in 30% sucrose (w/v) at 4°C overnight. Tissue was then embedded in OCT medium, snap-frozen, and stored at –80°C. Gut slices were rehydrated with PBS and blocked with 0.3% Triton X- in PBS and 10% of Normal Goat Serum for 1 h at room temperature. Samples were then incubated overnight with primary antibodies in 0.3% Triton X- in PBS and 1% of Normal Goat Serum. Samples were then washed with PBS and incubated for 1 h with phalloidin-iFluor-594 (1:500, Abcam) and with the appropriate secondary antibody (Invitrogen). After washing with PBS, samples were incubated for 5 minutes at room temperature with DAPI to stain nuclei (1:10000 in PBS). Images were acquired using a Leica TCS SP8 confocal microscope (Leica, Germany) equipped with a 63 × /1.4 numerical aperture oil-immersion objective and analyzed with ImageJ. For each mouse, 6–8 fields, each containing approximately 0.1 mm2 of epithelial-covered villus mucosa, were analyzed by a blinded observer. Number of positive stained cells per villi section per mm was calculated from at least 10 high power fields/section. For quantification of aggregated a-syn -specific signal, 8-10 pictures covering the whole gut section were taken using 10X objective, and a threshold was set where only the aggregate-specific signals were visible. The same threshold was applied to all images. The mean fluorescence intensity (MFI) of the selected aggregate signals was quantified using ImageJ software. Signals were normalized to the total surface area of each slice detected using the DAPI staining. Antibodies are listed in Supplementary Table 1.

### 2.6 *EX VIVO* CULTURES OF SPLEEN MACROPHAGES

Spleens were dissected from abdominal cavity and filtered through a 40-μm nylon strainer. Red cell lysis buffer was used to remove red cells. A single splenic cell suspension was obtained.^80^ Cells were cultured in Roswell Park Memorial Institute (RPMI) medium 1640 RPMI 1640 (BioConcept 1-41F01-I) supplemented with 10% FBS, 2mM L-Glutamine and antimicrobials (Penicillin-Streptomycin Pen 10’000 IU/ml Strep 10 mg/ml and amphotericin B (250 μg/ml BioConcept). Mouse microglia BV2 cells ^81^ from Elabscience (No.: EP-CL-0493) were cultured in parallel for each spleen culture preparation and served as controls. Spleen macrophages (SPMs) differentiate into the M1 phenotype after stimulation with LPS (100 ng/ml) ± IFN-γ (10 ng/ml).^82^ To monitor M1 status, of SPMs and BV2 cells, TNF-α (MTA00B) and IL-6 (M6000B) levels were determined using enzyme-linked immunosorbent assay (ELISA) kits (ELISA Development System; R&D Systems, McKinley Place, MN, USA) following the manufacturer’s protocol.^75^

### 2.7 LUMINEX-BASED MULTIPLEX CYTOKINE ANALYSIS

Cytokines from murine plasma samples, including IFN-γ, IL-1β, IL-2, IL-4, IL-6, IL-10, IL-17A, and TNF-α were measured simultaneously using the multiplexed kit “Mouse High Sensitivity T Cell Magnetic Bead Panel” (MHSTCMAG-70K, MILLIPLEX, EMD Millipore Corp., Billerica, MA, USA). The assay was performed according to the manufacturer’s instructions with an overnight incubation step (17 hours) at 4 °C. Samples were analyzed as single values and two controls with corresponding acceptance ranges, provided with the kit, were used for quality control purposes. The readout was performed on a FLEXMAP 3D® instrument (Luminex Corp., Austin, TX, USA). Data were acquired using Luminex xPONENT® software (version 4.3) and mean fluorescence intensity (MFI) was determined. Bio-Plex Manager™ software (version 6.2) (Bio-Rad, Hercules, CA, USA) was used for back-calculation of unknown cytokine concentrations.

### 2.8 GENE EXPRESSION ANALYSIS

Real-time quantitative PCR was performed using Taqman™Assay Reagents using the Step One Detection System (Applied Biosystems), according to manufactures protocol. The assay IDs are reported in *Supplemental Table 2*. Quantification of the abundance of target gene expression was determined relative to β-actin with respect to the control group by using the delta delta C_t_ (2^−ΔΔCt^) comparative method, the results expressed as arbitrary units (AU). Relative fold changes over WT are indicated.^71–74^

### 2.9 WESTERN BLOT ANALYSIS

Protein extracts were prepared as previously described.^83^ Tissues were homogenized in lysis buffer (0.33 M sucrose/8 mM Hepes, pH 7.4 and protease inhibitors) and quantified using the BCA protein determination method (Bio-Rad, Hercules, CA). Protein samples were diluted to equivalent volumes containing 20 μg of protein and boiled in an equal volume of Laemli SDS boiling buffer (Sigma) for 10 min. Samples were loaded into a 9-12% SDS-polyacrilamide gel and separated by electrophoresis for 3 h at 100 V. Proteins were transferred to polyvinylidene difluoride membrane (Amersham Biosciences, Piscataway, NJ) for 1.5 hr at 300 mA. After blocking of nonspecific binding with 5% non-fat dry milk in TBST, the membranes were probed with primary antibodies and processed as described (*Supplementary Table 1*). Densitometric analysis was performed using ImageQuantity One. Data were normalized to β-actin, values of phosphorylated GSK-3β (pTyr^216^ GSK-3β); phosphorylated α-syn (pSer^129^ α-syn) and phosphorylated tau (pSer^396^ tau) were normalized to total GSK-3β, α-syn, and tau, respectively, before statistical analysis of variance and values expressed as percent changes (%) of WT controls.^83^ Dashed lines (in white) indicate discontinuous bands (nonsequential lanes) taken from the same blot, at the same molecular weight (mass – kDa) in order to better represent the mean signal from all values (5-6 individual blots/genotype/treatment) for that particular group. Corresponding control bands (loading controls) match experimental bands.

### 2.10 STATISTICAL ANALYSES

Statistics was carried out using GraphPad Prism Software Version 9 (GraphPad Software, San Diego, California, USA). Significance was calculated using the unpaired, two-tailed Student’s t-test; Two-way or Three-way ANOVA, with Tukey’s multiple comparison or Bonferroni-post hoc test, as appropriate. Graphs displayed the mean ± SEM. *P <0.05, **P < 0.01 ***P < 0,001, ****P < 0.0001.

## 3. RESULTS

### 3.1 LRRK2 G2019S INTERACTS WITH AGEING AND CHRONIC LOW-GRADE INFLAMMATION TO IMPAIR NIGROSTRIATAL DOPAMINERGIC FUNCTION

To address the interaction between *LRRK2* G2019S, low grade chronic inflammation, and ageing, wild type (WT) and hemizygous C57BL/6J-TG (LRRK2*G2019S)2AMjff/J transgenic (G2019S) mice were exposed to intraperitoneal (ip) injection of a low dose (0,1 mg kg^−1^) of lipopolysaccharide (LPS, Escherichia coli serotype O111:B4), administrated twice a week for 12 consecutive weeks. As ageing is recognized as the primary risk factor influencing the clinical presentations and the progression of both sporadic and LRRK2 PD ^22,31^ treatments were performed in two different age groups, 3M (young adult) and 7M (at the beginning of middle-age) (*Supplementary Figure 1*). We first monitored motor coordination with the accelerated Rotarod. In NaCl-treated mice, a non-significant age-dependent motor deficit was observed in G2019S compared to WT mice (*Supplemental Figure 1C*). Both WT and G2019S mice treated with LPS showed a significant decrease in the Rotarod starting at 11 M, when compared to their motor performance at 3 M (Basal, -LPS). With ageing, however, only G2019S mice showed a stable significant deficit (*Supplemental Figure 1 C*). These results suggest a synergistic effect between *LRRK2* G2019S, ageing, and chronic low-grade systemic inflammation.

### 3.2 LRRK2 G2019S INTERACTS WITH AGEING AND CHRONIC LOW-GRADE INFLAMMATION TO PROMOTE NIGROSTRIATAL NEURODEGENERATION

Next, we performed a stereological quantification of DA neurons in the SNpc of NaCl- and LPS-treated mice of both genotypes at 3, 6, 10 and 16 M of age. Within the NaCl group, we did not find significant differences between G2019S WT mice from 3 to 16 M; however, within genotypes, a significant decrease in the total number of TH^+^ neurons was observed by 16 M in both WT and G2019S NaCl-treated mice (**Figure 1 A, C**). The chronic low-dose LPS regimen in 3 M old mice for 12 consecutive weeks did not affect the total number of TH^+^ neurons by 6 M of age in both genotypes (**Figure 1 B, C**), whereas starting LPS treatment at middle age (7 M) for 12 weeks until 10 M, significantly reduced TH^+^ neuron numbers in G2019S (÷ 70% loss) compared to WT counterparts (÷ 30% loss). After this time-point, TH^+^ neurons further declined until 16 M in TG and WT mice, with TH^+^ neuron loss being significantly greater in G2019S (÷ 80%) compared to WT (÷ 40%) (**Figure 1 A, B, C**). Quantitative confocal laser microscopy on striatal sections revealed an age-dependant reduction of striatal TH-immunofluorescent (IF) reaction in both genotypes, without significant differences between WT and G2019S genotypes (**Figure 1A, C**). Starting the LPS regimen at 3 M did not affect TH-IF at 6 M, in both genotypes, whereas starting the LPS chronic regimen at middle-age significantly reduced striatal TH-IF reaction (**Figure 1 B, C**). Within the LPS-treated group, striatal TH immunoreactivity was significantly reduced in G2019S compared to WT mice. (**Figure 1 B, C**).

**Figure 1.**
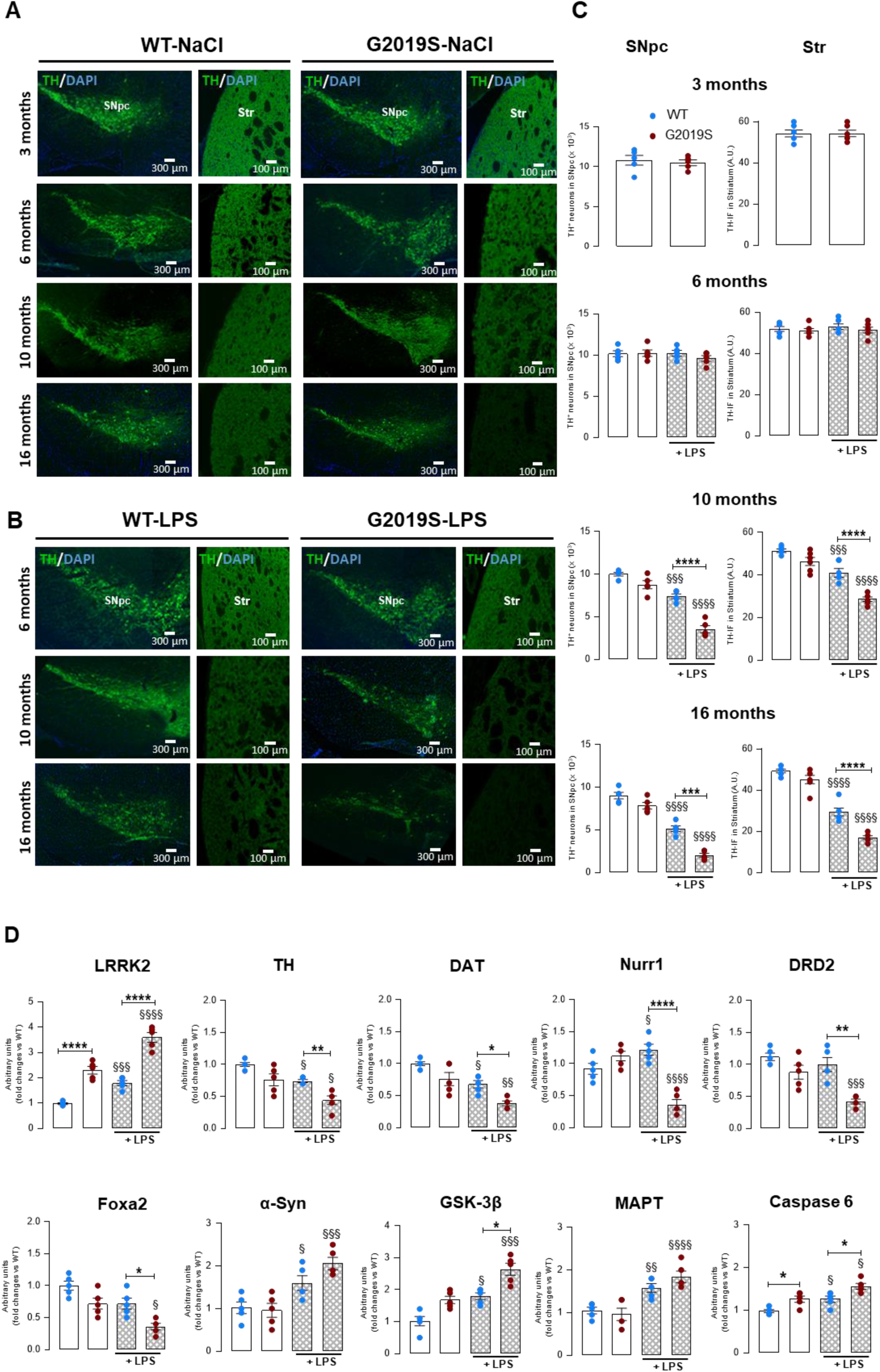
G2019S interacts with ageing and chronic low-grade inflammation to drive nigrostriatal dopaminergic neurodegeneration. **A**. Representative confocal images of the SNpc and the Str of WT and G2019S mice from 3- to 16 months (M) of age under NaCl (**A**) or a low dose of LPS (0.1 mg Kg^−1^, twice weekly, i.p.) administrated for 12 consecutive weeks (**B**). Tissues were immunostained with TH (green) and the nuclear marker DAPI (blue). Scale bars, SNpc, 300 μm; Str, 100 μm. **C**. Total number of TH^+^ neurons (mean ± SEM, 5-6 mice/age-group/treatment/genotype) counted in the right and left SNpc and striatal TH-IF at 3, 6, 10 and 16 M under NaCl or LPS exposure. A significant decrease in the total number of TH^+^ neurons was measured by 16 M in both WT (§ < 0.05 *vs* WT 3 M) and G2019S mice (§§P < 0.01 *vs* TG 3 M). G2019S, low-grade inflammation and ageing robustly reduced TH^+^ neuron numbers and Str-IF in G2019S vs WT mice (P< 0.0001). **D**. Expression levels of the indicated genes measured in the SNpc of WT and G2019S mice treated with NaCl or LPS at 10 M of age. Data were normalized over WT untreated and expressed as mean ± SEM; n: 5-6 mice/age-group/treatment/genotype; Significance analyzed by ANOVA followed Tukey’s *\*P*< 0.05, ** *P* < 0.01, *** P < 0.001; **** P < 0.0001 *vs* WT within the same treatment group; §*P*< 0.05, §§ *P* < 0.01, §§§ P < 0.001; §§§§ P < 0.0001 *vs* NaCl within each genotype.

### 3.3 LRRK2 G2019S MICE DISPLAY DOWNREGULATION OF DOPAMINERGIC NEURON TRANSCRIPTS, INCREASED PHOSPHORYLATED α-SYN, GSK-3β and TAU

Next, we focused on the 10 M group to assess gene and protein levels in the ventral midbrain (VM) of WT and G2019S mice (**Figures 1D and Figure 2 A-C**). Within the NaCl-treated group, *LRRK2* gene expression was significantly up-regulated in G2019S compared to WT mice (**Figure 1 D**). LPS exposure increased *LRRK2* expression in both genotypes, with a more significant impact in G2019S mice (**Figure 1 D**). DA neuron mRNA species (*Th, Slc6a3* (DAT), *Drd2, Nr4a2* (NURR1), *Foxa2*) were significantly down-regulated upon LPS exposure in G2019S compared to LPS-treated WT mice (**Figure 1 D**). LRRK2 regulates tau phosphorylation through activation of glycogen synthase kinase-3β (GSK-3β),^84–86^ which, in turn, is involved in mDAn death.^72,87–91^ LPS treatment significantly increased GSK-3β transcript levels both in WT and G2019S mice, with a significant higher impact in G2019S mice (**Figure 1 D**). Furthermore, LPS induced a significant increase in *SNCA* (α-syn) and *MAPT* (microtuble associated protein tau) gene expression levels in both WT and G2019S mice, with significant higher levels detected in G2019S compared to WT mice (**Figure 1 D**). Finally, the cell death marker Caspase 6 was up-regulated in G2019S compared to WT mice in both NaCl and LPS groups (**Figure 1 D**). Immunofluorescent staining and Western blot analyses confirmed these data (**Figure 2 A-C**). We found increased LRRK2 levels in SNpc DAT^+^ neurons of NaCl-treated G2019S mice compared to WT counterparts (**Figure 2 A**). Additionally, low-grade inflammation increased LRRK2 levels in both WT and G2019S mice, with a more significant effect in G2019S mice (**Figure 2 A**). Upon LPS treatment, both α-syn and α-syn phosphorylated at Ser-129, which is the dominant pathological modification of α-syn in both familial and sporadic PD ^91,92^ were increased in SNpc DAT^+^ neurons of G2019S mice compared to WT mice of the same age and treatment group (**Figure 2 B**). Likewise, both proteins were up-regulated in G2019S compared to WT counterparts, as determined by WB (**Figure 2 C**). Importantly, TH and Nurr1 protein levels were significantly reduced in in G2019S compared to WT SNpc, whereas active pTyr^216^-GSK-3β protein levels determined by WB analysis were significantly increased in NaCl- and LPS-treated G2019S compared to WT mice (**Figure 2 C**). LPS treatment increased tau phosphorylation at Ser396, ^16,84–88,90,91^ when compared to WT counterparts (**Figure 2 C**). These data suggest that LRRK2 G2019S synergizes with ageing and low-grade chronic inflammation to promote the up-regulation of several key factors involved in mDAn dysfunction/death.

**Figure 2.**
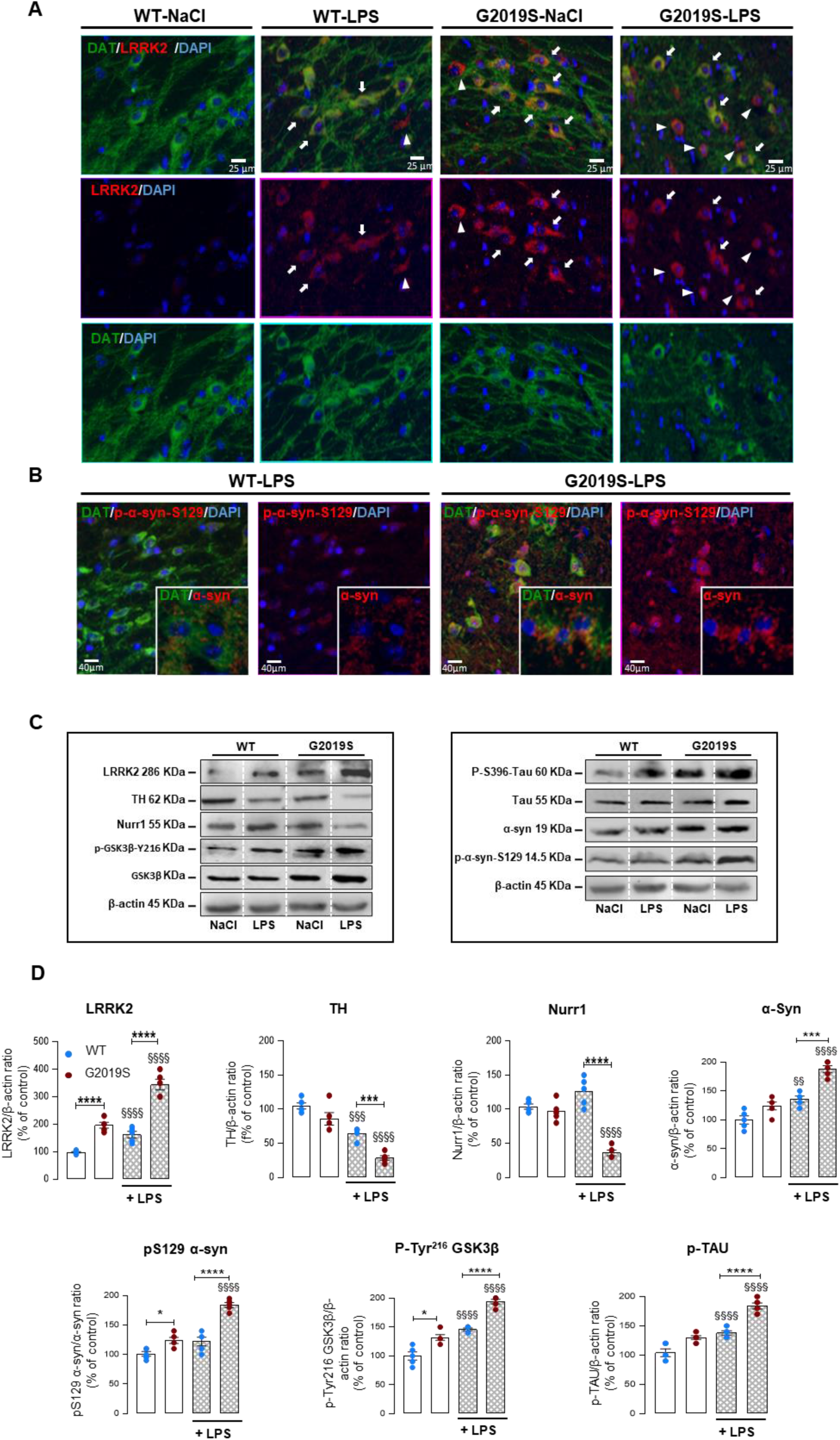
Low-grade inflammation induces LRRK2, active pGSK-3β, α-syn, and Tau levels in aged G2019S VM. **A**. Triple immunofluorescent staining of midbrain sections with LRRK2 (red), the dopamine transporter (DAT, green), and DAPI (blue), showing increased LRRK2-IF reaction colocalizing in SNpc DAT^+^ neurons of 10 M-old NaCl G2019S mice, *vs* WT counterparts. LPS further magnified LRRK2-IF localized to both DAT-positive neurons (arrows) and DAT-negative neurons (arrowheads) in G2019S mutant mice. Scale bars, 25 μm. **B**. Triple immunofluorescent staining with DAT (green), phosphorylated (p) α-syn (in red) and DAPI (blue), and α-syn (boxed) showing a greater pα-syn and α-syn IF signals in SNpc DAT^+^ and some DAT^−^ neurons of 10 M-old G2019S- *vs* WT-LPS. Scale bars, 40 μm. **C**. Representative Western blots showing proteins levels of LRRK2, TH, Nurr1, pTyr^216^-GSK-3β, GSK-3β, P-S396-Tau, Tau, α-syn, p-and α-syn-S129 in 10 M-old WT and G2019 mice under NaCl or LPS treatment. Quantification of protein levels relative to the loading control is shown, values of phosphorylated GSK-3β (pTyr^216^ GSK-3β); phosphorylated α-syn (pSer^129^ α-syn) and phosphorylated tau (pSer^396^ tau) were normalized for each respective control (total GSK-3β, α-syn, and tau, respectively). Data represent the mean % ± SEM of 5-6 mice/age-group/treatment/genotype. Statistical significance analyzed by ANOVA followed Tukey’s \**P*< 0.05, ** *P* < 0.01, *** P < 0.001; **** P < 0.0001 *vs* WT within the same treatment group; §*P*< 0.05, §§ *P* < 0.01, §§§ P < 0.001; §§§§ P < 0.0001 *vs* NaCl within each genotype.

### 3.4 LRRK2 G2019S SYNERGYSES WITH AGEING AND SYSTEMIC LOW-GRADE INFLAMMATION TO EXACERBATE ASTRO- AND MICROGLIOSIS

Next, we examined inflammatory reactions in the SNpc of WT and G2019S mice. Coronal SNpc and Str sections were immunostained for the astrocytic cell marker, GFAP, the macrophage/microglial cell marker, IBA1, the chemokine receptor, CCR2, which is known to be expressed on peripheral monocytes and macrophages but not in brain microglia, ^93–95^ and the T-cell marker CD3. We found that G2019S mice displayed an age-dependant increase of GFAP^+^ astrocytes starting at 10 M, whereas in WT mice GFAP^+^ astrocytes significantly increased by 16 M compared to WT mice at 3M (**Figure 3 A**). Furthermore, at 10 and 16 M, LPS further enhanced astrocyte cell numbers in G2019S mice (**Figure 3A**). The middle-age group was the most vulnerable group in both genotypes; here, LPS-treated G2019S mice show the most significant increase in GFAP^+^ astrocytes (**Figure 3 B-C**). At 10 M, LPS-treated G2019S mice showed a significant astrocytic hypertrophy compared to WT counterparts (**Figure 3 C, D**). In line with these data, GFAP was up-regulated at the mRNA and protein levels in the VM of NaCl and LPS-treated G2019S mice (**Figure 3 E, F**). Furthermore, we observed neurite loss and TH^+^ neuron atrophy associated to poor contact with GFAP^+^ astrocytes in the SNpc of 10 M-old LPS-treated G2019S mice (**Figure 3 D**). On the contrary, WT mice under LPS displayed longer TH^+^ neurites, and TH neuronal cell showed closed interactions with GFAP^+^ astrocytic processes, similar to what observed in NaCl-treated WT mice (**Figure 3 C**). In the striatum, we detected a significant increase of GFAP^+^/DAPI^+^ astrocytes both at 10 and 16 M in G2019S mice compared to WT mice, which was enhanced by LPS chronic exposure (*Supplementary Figure 2 A-B*).

**Figure 3.**
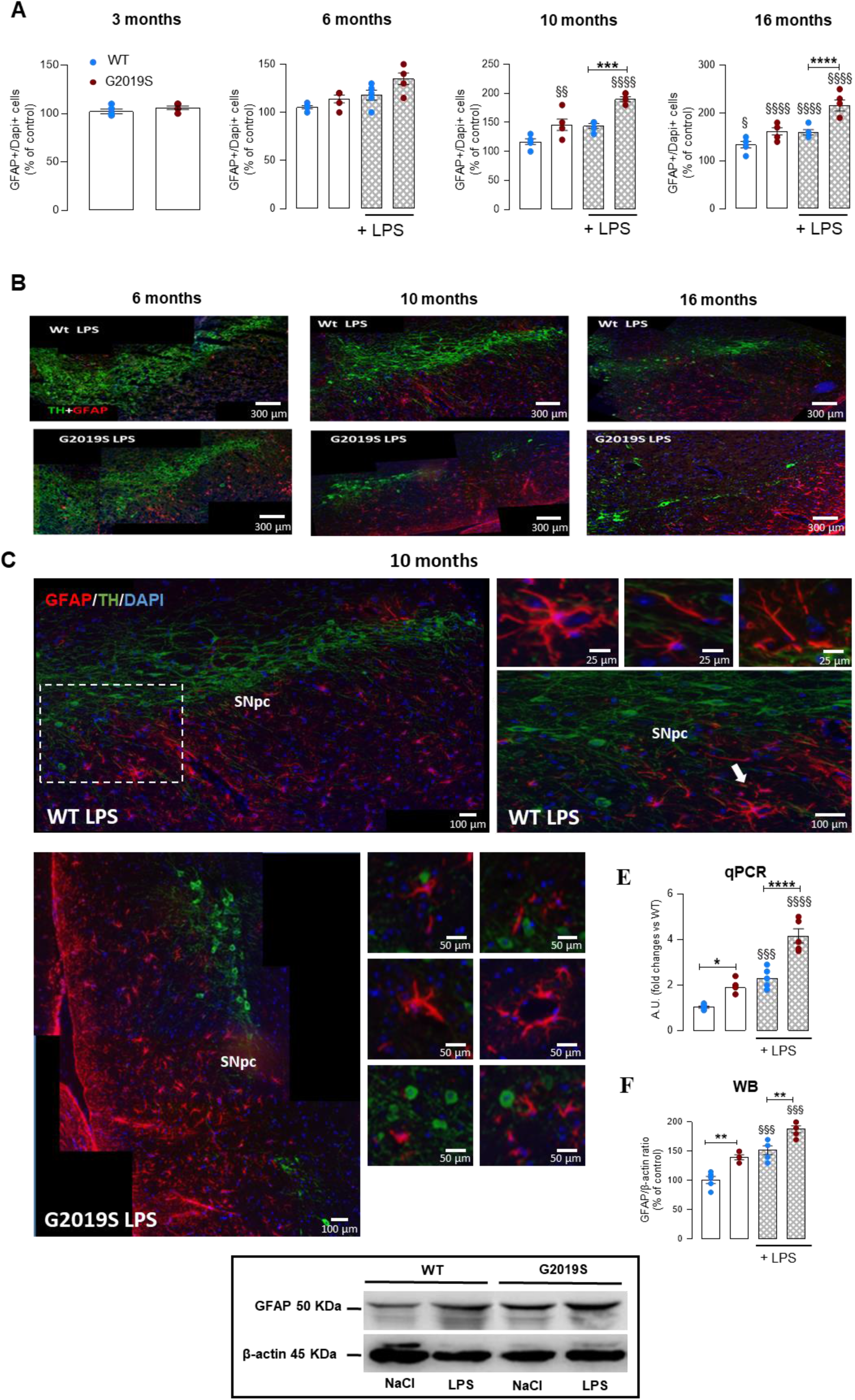
LRRK2 G2019S combined with low-grade inflammation and ageing exacerbates astrocyte hypertrophy in the VM. **A**. Quantification of GFAP^+^/Dapi^+^ astrocytes in SNpc of 3 to 16 M WT and G2019S mice under NaCl/LPS regimen. No changes in GFAP^+^/Dapi^+^ astrocytes under NaCl in both genotypes were observed until 6 M, whereas from 10 M on, NaCl-treated G2019S mice displayed a greater number of astrocytes (P ≤ 0.01) *vs* the 3 M age group, and much greater *vs* NaCl-treated WT counterparts (P < 0.05). G2019S, ageing, and LPS further increase (P ≤ 0.001) cell number of G2019S *vs* WT by 16 M. **B**. Representative confocal images of triple immunofluorescent staining with TH (green), GFAP (red) and DAPI (blue) in SNpc at 6, 10 and 16 M, under LPS, showing enhanced astrocytosis and reduced TH^+^ neurons at 10 and 16 M in G2019S *vs* WT counterparts. Scale bars, 300 μm. **C-D**. Representative confocal images of triple TH^+^/GFAP^+^/Dapi^+^ cells in SNpc of WT (**C**, Scale bars 100 μm, boxed images, 25 μm) and G2019S (**D**, Scale bars 100 μm, boxed images, 50 μm) mice under LPS, showing increased TH^+^ neurite extension and healthier TH neuronal cell bodies with closed interactions with GFAP^+^ astrocyte processes of WT (**C**) *vs* G2019S (detailed in text) (**D**). **E-F:** qRT-PCR and WB analysis showing higher expression of GFAP transcript and protein levels in G2019S mice both under NaCl and LPS regimen *vs* WT counterparts. Data (GFAP/β-actin ratio relative to controls), represent the mean % ± SEM of 5-6 mice/age-group/treatment/genotype. Significance analyzed by ANOVA followed Tukey’s *\*P*< 0.05, ** *P* < 0.01, *** P < 0.001; **** P < 0.0001 *vs* WT within the same treatment group; §*P*< 0.05, §§ *P* < 0.01, §§§ P < 0.001; §§§§ P < 0.0001 *vs* NaCl within each genotype.

Similarly, the number of IBA1^+^ microglial cells increased as a function of age and LPS exposure, with significant higher numbers in G2019S mice compared to WT mice both in the SNpc and Str (**Figure 4 A, B, C** and *Supplementary Figure 2 C-D*). Microglial cells shifted to a reactive phenotype with age and LPS treatment in both genotypes, characterized by increased cell body area with less ramified processes (**Figure 4 B-C**). G2019S mice under LPS exposure showed significantly higher levels of major inflammatory transcripts including Macrophage antigen complex-1 (*Mac1*), nuclear factor kappa-light-chain-enhancer of activated B cell (*NF-κB*), inducible nitric oxide synthase (*NOS2*), phagocyte oxidase (*gp91PHOX*), a major NADPH oxidative stress transcript member, ^96^ cyclooxygenase 2 (*COX2, Supplementary Figure 3*), the cytokines *TNF-α, TNF-R, IL-18*, which are inflammatory cytokines induced by *NF-κB* activation, ^97^(**Figure 4 D and *Supplementary Figure 3***), and the C-X-C motif chemokine 10, *CXCL10* (**Figure 4 D**) promoting leukocyte recruitment and neuronal injury.^98^ On the contrary, the C-C chemokine, *CCL3* significantly decreased under LPS treatment in G2019S VM (*Supplementary Figure 3*). Similarly, in WT mice of the same age- and treatment-group, such inflammatory markers increased under low-grade inflammation, albeit levels were lower compared to G2019S mice (**Figure 4 D**, and *Supplementary Figure 3*). While we did not find differences in *IL-1β* mRNA levels, at the protein level, pro-caspase-1 promoting caspase-1 activation, ^99^ was significantly elevated in G2019S VM. These data were supported by higher levels of mature IL-1β protein in G2019S VM (**Figure 4 E**). The cytokines *IL 4* and *IL 10* were undetectable in both WT and G2019S mice under LPS regimen in the same age group. In line with an exacerbated pro-inflammatory status of G2019S VM, we found increased IBA-1, NF-κB-65, iNOS and gp91Phox protein levels by Wb analysis, as compared to WT LPS-treated counterparts (**Figure 4 E**).

**Figure 4.**
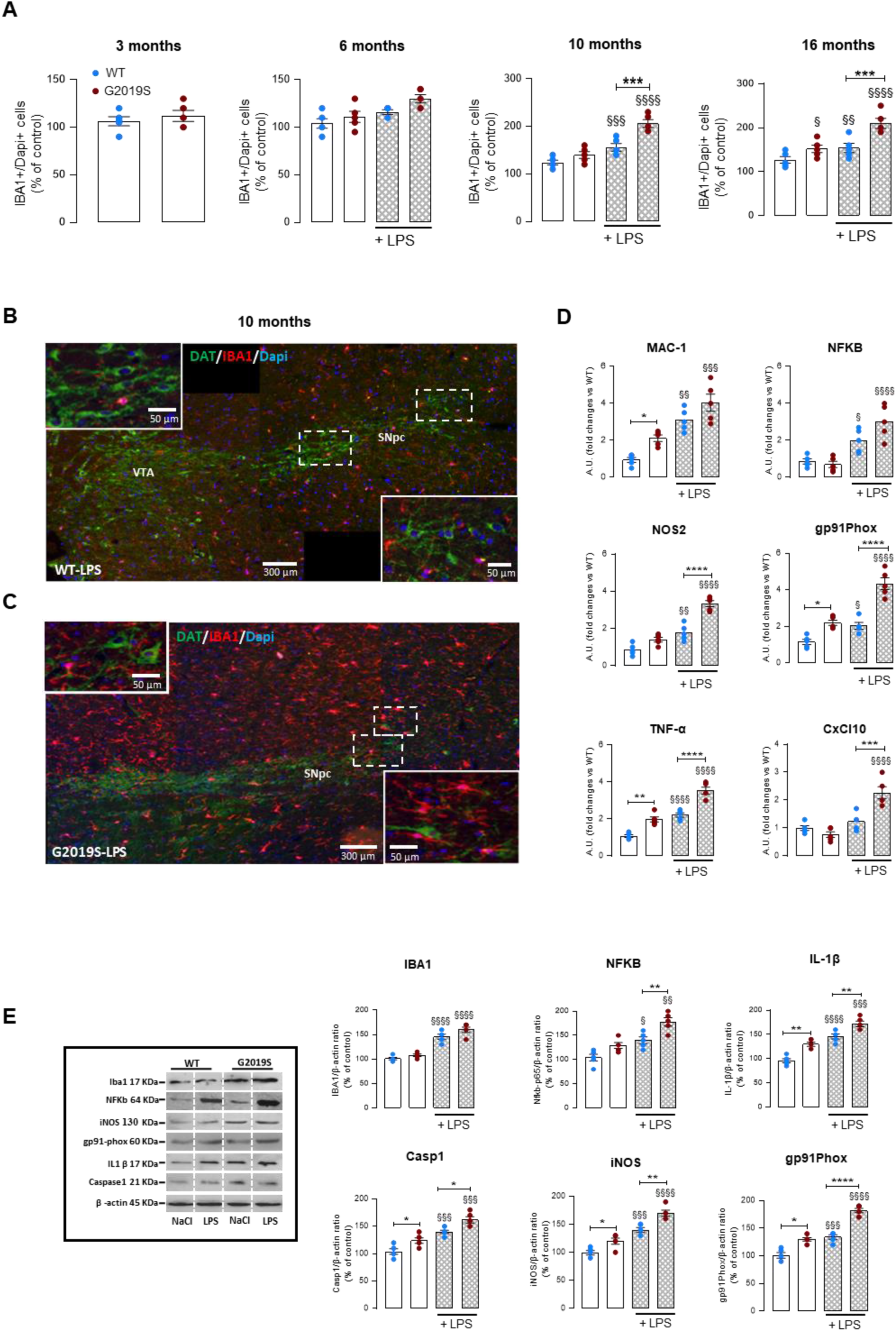
LRRK2 G2019S, low-grade inflammation, and ageing exacerbate microglia inflammatory reactions in the VM. **A**. Quantification of IBA1^+^/Dapi^+^ cells in SNpc sections of WT and G2019S mice upon LPS or NaCl treatment at the indicated timepoints. **B-C**. Representative confocal images of triple immunofluorescent staining for DAT (green), IBA1 (red), and DAPI (blue) in SNpc sections of WT and G2019S mice upon LPS or NaCl treatment at the indicated timepoints. Scale bars; 300 μm, boxed areas 50 μm. **D**. qRT-PCR in SNpc tissues performed at 10 M of age showing upregulation of major inflammatory and pro-oxidant transcripts in G2019S mice. **E**. Representative western blots of IBA1, NF-κB, iNOs, gp91PHOX, IL-1β protein, and caspase-1 in G2019S VM, as compared to WT counterparts. Quantification of protein levels relative to the loading control is shown. Mean ± SEM of 5-6 mice/age-group/treatment/genotype. Significances analysed by ANOVA followed Tukey’s \**P*< 0.05, ** *P* < 0.01, *** P < 0.001; **** P < 0.0001 *vs* WT within the same treatment group; §*P*< 0.05, §§ *P* < 0.01, §§§ P < 0.001; §§§§ P < 0.0001 *vs* NaCl within each genotype.

### 3.5 LRRK2 G2019S PROMOTES THE RECRUITMENT OF PERIPHERAL MONOCYTES AND T-CELLS

Next, we examined whether LRRK2 G2019S alters peripheral immune cell infiltration. One of the key mechanisms involved in monocyte recruitment is the CCL2/CCR2 signaling, which plays a key role in the recruitment of peripheral monocytes into the SN in a PD model of α-syn-induced inflammation and neurodegeneration.^100^ Using triple immunofluorescent staining, we monitored macrophage/monocyte trafficking from 3 M to 16 M. Undetectable or low CCR2-IF signal was found in 3 to 6 M old mice in both NaCl- or LPS-treated mice, irrespective of the mouse genotype (**Figure 5A, B**). Starting at 10 M, we found significant higher numbers of CCR2^+^/IBA1^−^ cells in the midbrain of LPS-treated G2019S mice compared to WT mice (**Figure 5 A, C**). Remarkably, reactive GFAP^+^ astrocytes (*Supplementary Figure 4*) and IBA1^+^ microglia (**Figure 5 C**) were observed closed to infiltrating CCR2^+^ macrophages/monocytes. These results were confirmed by WB analysis that showed a significant increase of CCR2 protein levels in the VM of 10 M-old G2019S mice, in both NaCl and LPS experimental groups (*Supplementary Figure 4*). Furthermore, immunofluorescent staining revealed CD3^+^ T cell brain infiltration in the SNpc of 10 M-old LPS-treated G2019S mice compared to WT mice (**Figure 5 E**). At the peripheral level, we found a greater spleen infiltration of CD3^+^ /LRRK2^+^ cells starting at 6 M in LPS-treated G2019S mice compared to WT spleens of the same age and treatment (**Figure 5 F-G**). Increased LRRK2 protein was found in the spleen of G2019S mice, both under NaCl and LPS treatment by 6 M (**Figure 5 F-G**). To examine the response of spleen macrophages (SPMs) as a function of age x genotype x LPS regimen, *ex vivo* individual cultures of SPMs established from WT and G2019S mice of different age-groups were differentiated into the M1 phenotype after stimulation with LPS (100 ng/ml) ± IFN-γ (10 ng/ml) (**Figure 6A**). To monitor M1 status, we evaluated TNF-α and IL-6 release in the conditioned medium (CM). First, we assessed the response at basal conditions, in untreated individual mice of different ages and found no difference between G2019S and WT SPMs in the release of TNF-α and IL-6 from 6 to 16 M of age. LPS treatment promoted a significant TNF-α and IL-6 release in both primary splenocytes and BV2 cells, used as control (**Figure 6 B-C**). M1-polarized SPMs from LPS-treated G2019S mice expressed higher amounts of both cytokines starting at 6 M (**Figure 6 D**). Moreover, LRRK2-G2019S synergized with LPS and IFN-γ to promote TNF-α and IL-6 release in G2019S mice compared to WT SPMs derived from WT counterparts of the same age and treatment (**Figure 6 D**). These data indicate that in the spleen, G2019S and low-grade inflammation shift SPMs towards a proinflammatory phenotype starting at 6 M. Next, we measured the protein level of cytokines in the serum of WT and G2019S mice in response to LPS at different time points. At the baseline (3 M), LRRK2 G2019S mice showed increased levels of IL-6 and IL-4 compared to WT mice (Figure 7). Furthermore, we found a significant increase in the levels of IL-6, IL1-beta, IFN-γ, IL-2, TNF-α, IL-17A, and IL-4 in 6 M old G2019S LPS-treated mice compared to WT mice. At 16 M, G2019S LPS-treated mice showed increased levels of IL-6, IFN-γ, TNF-α compared to WT mice (Figure 7A).

**Figure 5.**
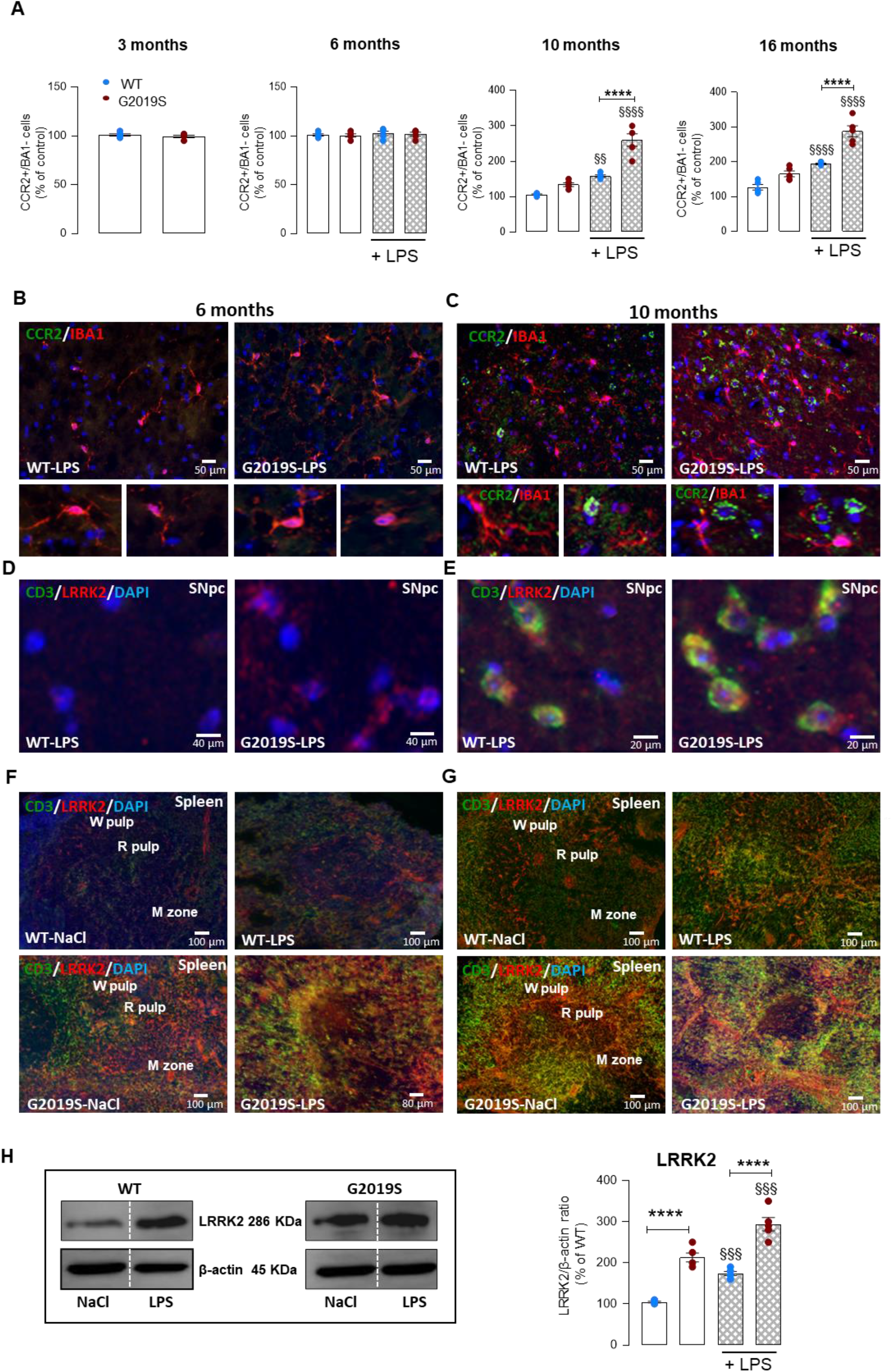
LRRK2 G2019S synergizes with low-grade inflammation to promote macrophage and CD3^+^ T cell recruitment to the brain in aged mice. **A**. Quantification of CCR2^+^/IBA1^−^ cells in the SNpc in 3 to 16 M-old WT and G2019S mice under NaCl/LPS. **B,C**. Triple immunostaining of IBA1 (red), CCR2 (green), and DAPI (blue) at 6 M (**B**, Scale bar 50 μm) and 10 M (**C**, Scale bar, 50 μm) in WT and G2019S mice under LPS. **D-E**: Representative immunostaining of CD3 T-cell marker (green), LRRK2 (red), and DAPI (blue) in the SNpc of 6 M (**D**, Scale bar, 40 μm) and 10 M (**E**, Scale bar, 20 μm) WT and G2019S mice under LPS. **F,G**. Immunostaining of CD3^+^ cells expressing LRRK2 in spleen sections of 6 M and 10 M-old WT and G2019S mice under NaCl/LPS (Scale bar, 100 μm). **H**. Representative Western blot of LRRK2 in the spleen of WT and G2019S mice under NaCl/LPS at 6 M showing significantly up-regulated LRRK2 protein level in G2019S vs WT counterparts. Quantification of protein levels relative to the loading control is shown. Mean ± SEM of 5-6 mice/age-group/treatment/genotype. Significances analysed by ANOVA followed Tukey’s \**P*< 0.05, ** *P* < 0.01, *** P < 0.001; **** P < 0.0001 *vs* WT within the same treatment group; §*P*< 0.05, §§ *P* < 0.01, §§§ P < 0.001; §§§§ P < 0.0001 *vs* NaCl within each genotype.

**Figure 6.**
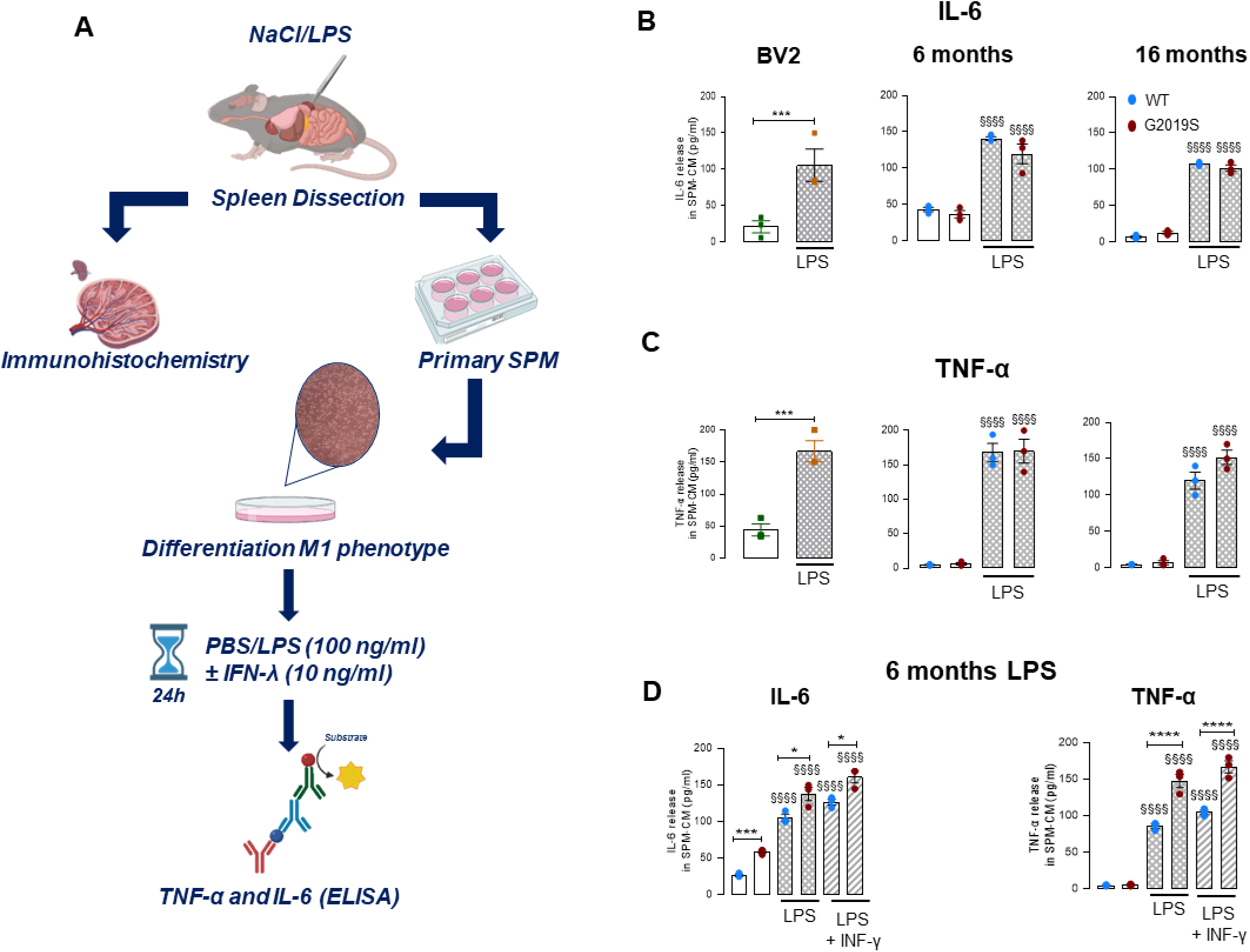
LRRK2 G2019S and low-grade inflammation exacerbate the response of spleen macrophages to LPS and IFN-γ. **A**. Schematic illustration of the experimental protocol performed in WT and G2019S mice treated with NaCl or LPS for 12 weeks. *Ex vivo* individual primary cultures of spleen macrophages (SPMs) established from WT and G2019S mice of different age-groups were differentiated into the M1 phenotype after stimulation with LPS (100 ng/ml) ± IFN-γ (10 ng/ml). **B-C**. The response of SPMs to LPS at 6 and 16 M was assessed in untreated individual mice. BV2 microglial cells were used as controls. TNF-α and IL-6 release measured in the conditioned media (CM) indicated no difference between G2019S and WT SPMs in the release of TNF-α and IL-6 from 6 to 16 M of age. **D**. M1-polarized SPMs from LPS-treated G2019S mice expressed higher amounts of TNF-α and IL-6 starting at 6 M. Note that IL-6 is significantly higher in G2019S *vs* WT, both before (*P* < 0.001) and after LPS. LRRK2 G2019S synergized with LPS and IFN-γ to promote higher TNF-α and IL-6 release in G2019S mice compared to WT SPMs derived from WT counterparts of the same age and treatment. Mean ± SEM of 5-6 mice/age-group/treatment/genotype. Significances analysed by ANOVA followed Tukey’s *\*P*< 0.05, *** *P* < 0.001; **** *P* < 0.0001 *vs* WT within the same treatment group; §§§§ *P* < 0.0001 *vs* NaCl within each genotype.

**Figure 7.**
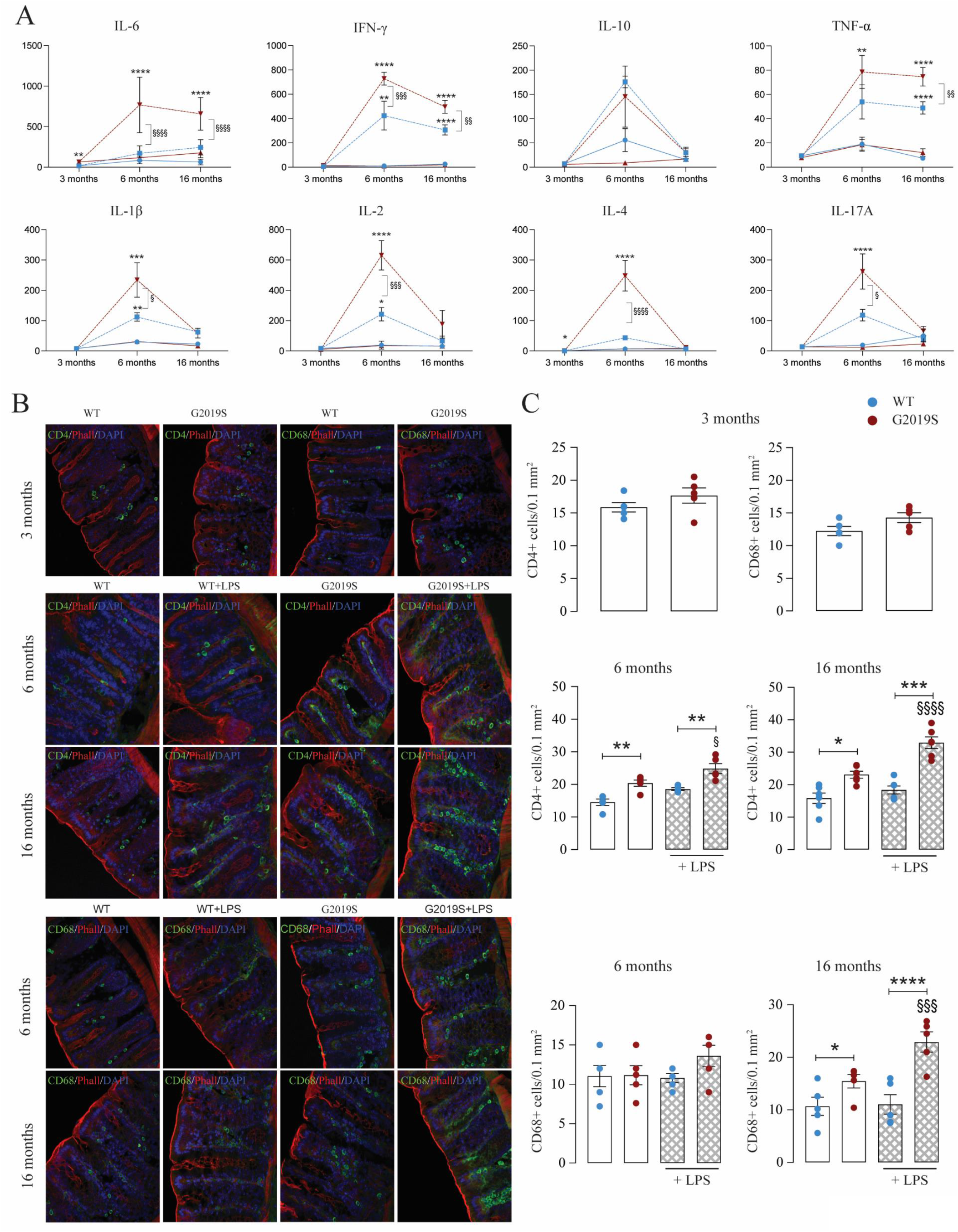
(**A**) Cytokines profile in the serum of WT and LRRK2 G2019S mice at the baseline (3 M) and 6-16 M. Mean±SEM, n=6. §§§§, p <0.0001; §§§, p <0.001, §§, p <0.001, §, p <0.05, LPS-treated vs NaCl-treated; **** p <0.0001, *** p <0.001, ** p <0.01, * p <0.05, LRRK2 G2019S vs LRRK2 WT, vs group indicated on the graph, two-way ANOVA with Bonferroni post hoc test; n=5 independent experiments. Data are expressed as pg/mL. (**B**) Intestinal cryosections stained against CD4 or CD68 (green), phalloidin (red), and DAPI (blue) from WT and LRRK2 G2019S mice treated with LPS or NaCl at the indicated time-points. Scale bar = 100 μm. (**C**) Analysis of CD4+ and CD68+ cells within colonic sections. Mean ± SEM; §§§§, p <0.0001; §§§, p <0.001, LPS-treated vs NaCl-treated controls; **** p <0.0001, *** p <0.001, ** p <0.01, * p <0.05, LRRK2 G2019S vs WT mice, two-way ANOVA with Bonferroni post hoc test; n=6. Cells counted from at least 10 high power fields (100X)/mouse.

### 3.6 GUT INFLAMMATION AND A-SYN AGGREGATION PRECEDE NEURODEGENERATION

Multiple lines of evidence support a role for gastrointestinal (GI) dysfunction and intestinal α-syn pathological accumulation in PD pathogenesis.^101^ A link between GI inflammation and neurodegeneration has also been proposed. However, whether intestinal immune cell activation precede neurodegenerative processes has not been investigated. To address this question and examine whether LRRK2 G2019S modulates intestinal immune cell infiltration, we assessed CD4^+^ T-cell and macrophage numbers in colonic tissues of WT and G2019S mice at different timepoints (3 M, 6M, and 16M). We observed a significant increase of CD4^+^ T-cells in the colon of untreated LRRK2 G2019S mice, compared to WT mice starting at 6 months of age (Figure 7B, C). LPS treatment significantly increased CD4^+^ T-cell infiltration in the colon of LRRK2 G2019S mice at 6 M, whereas no significant effect was observed in LRRK2 WT mice (**Figure 7B, C**). Furthermore, a significant increase of CD68^+^ macrophages was observed in 16 M-old LRRK2 G2019S compared to LRRK2 WT mice (**Figure 7B, C**). At 16 M, treated LRRK2 G2019S mice showed a significant increase in colonic CD68^+^ cells compared to LRRK2 WT mice (**Figure 7B, C**). Immunohistochemical analysis of colonic tissues with the aggregate specific α-syn antibody, MJF-14, showed the induction of aggregation of α-syn by LPS treatment in both 6 and 16 M-old LRRK2 G2019S and LRRK2 WT mice, with a significant stronger effect in LRRK2 G2019 mice (**Figure 8A, B**). However, neither the LRRK2 genotype nor the treatment affected the number of HuC/HuD positive neurons at the different timepoints of analysis (**Figure 9A, B**). These results suggest that the LRRK2 genotype is associated with age-dependent gut immune cell infiltration, which, combined with a low dose chronic inflammatory stimulus, may promote α-syn aggregation.

**Figure 8.**
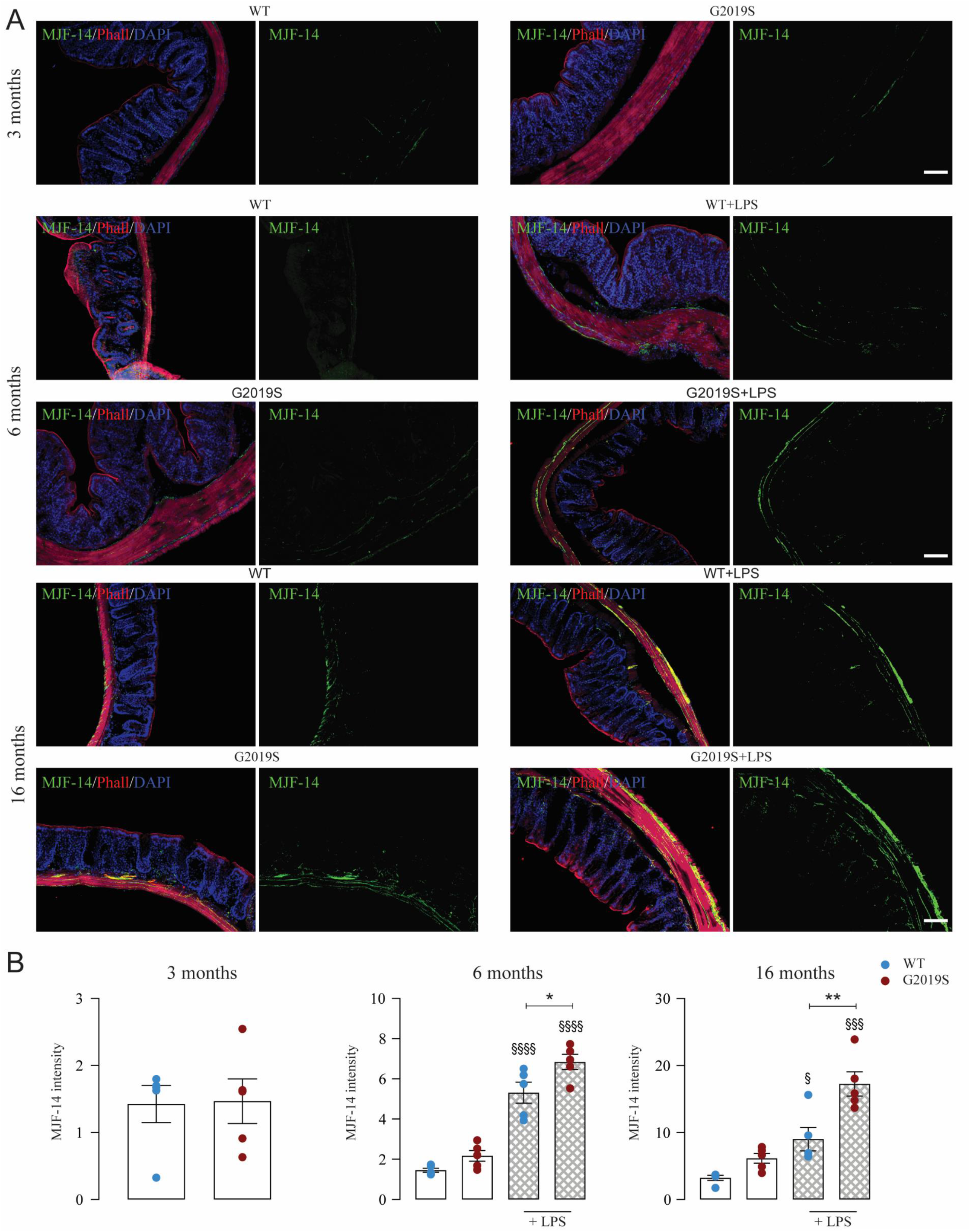
(**A**) Intestinal cryosections stained for aggregated α-syn (MJF-14, green), TH (red), phalloidin (magenta), and DAPI (blue) from WT and G2019S mice treated with LPS or NaCl at the indicated time-points. Scale bar = 100 μm. (**B**) Quantification of MJF-14 mean intensity within colonic sections. Mean ± SEM; §§§§, p <0.0001; §§§, p <0.00, LPS-treated vs NaCl -treated controls; **** p <0.0001, *** p <0.001, ** p <0.01, * p <0.05, LRRK2 G2019S vs WT mice, two-way ANOVA with Bonferroni post hoc test. Data were acquired from at least 10 high power fields (100X)/mouse, n=6.

**Figure 9.**
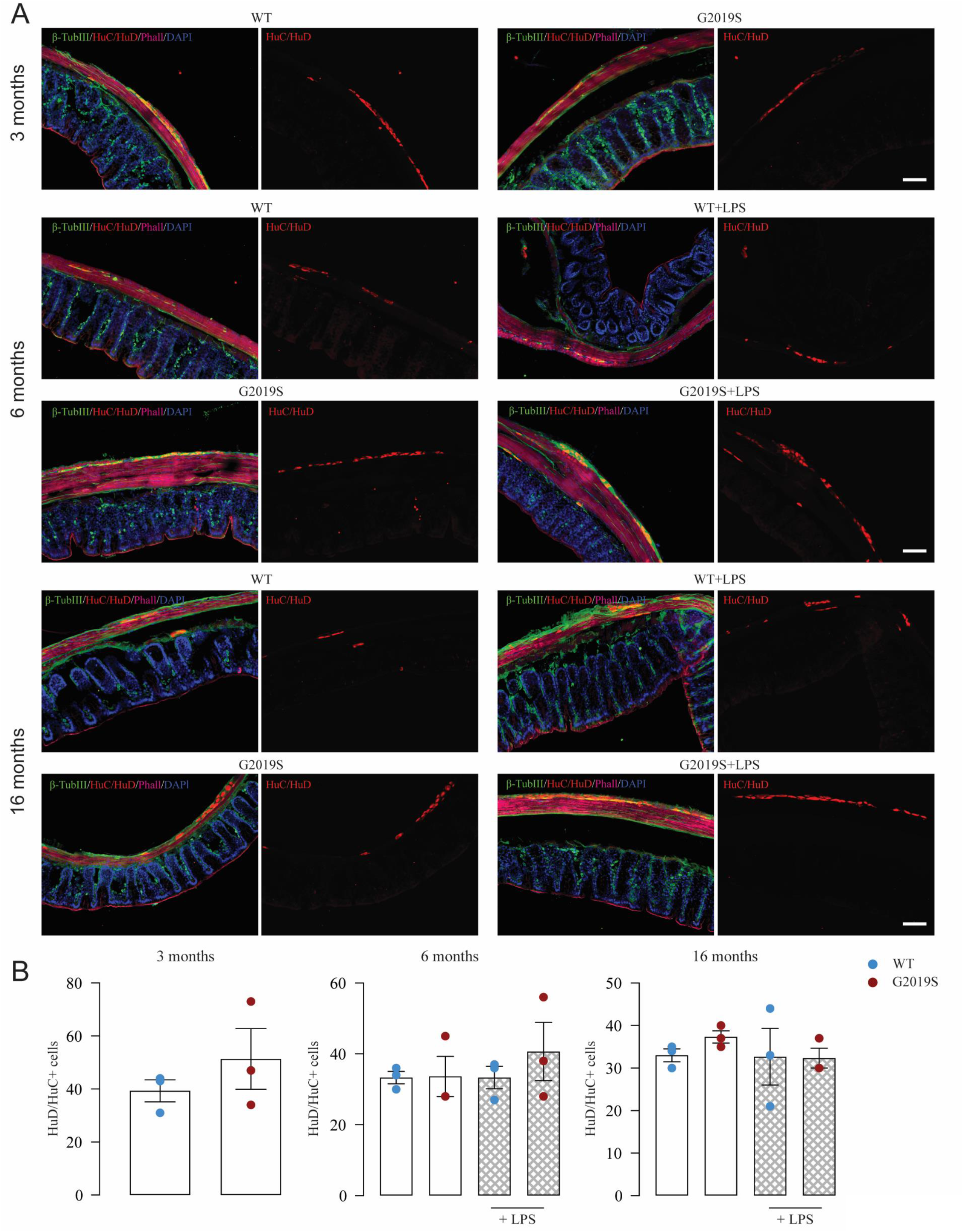
(**A**) Intestinal cryosections stained against HuD+HuC (red), ß-TubIII (green), phalloidin (magenta), and DAPI (blue) from WT and G2019S mice treated with LPS or NaCl at the indicated time-points. Scale bar = 100 μm. (**B**) Quantification of HuD+HuC positive cells within colonic sections. Mean ± SEM; §§§§, p <0.0001; §§§, p <0.00, LPS-treated vs NaCl -treated controls; **** p <0.0001, *** p <0.001, ** p <0.01, * p <0.05, LRRK2vs WT mice, two-way ANOVA with Bonferroni post hoc test. Data were acquired from at least 10 high power fields (100X)/mouse, n=6.

## 4 DISCUSSION

Here, we examined the role of the PD-associated *LRRK2* G2019S mutation in modulating central and peripheral immune responses as well as nigrostriatal DA vulnerability with ageing. We show that low-dose inflammation triggers mDAn degeneration in LRRK2 G2019S mice, macrophage/monocyte CNS infiltration as well as astro- and microgliosis. Interestingly, neurodegeneration is preceded by peripheral immune activation as well as increased CD4^+^ T-cell infiltration and α-syn aggregation in the colon. Furthermore, we show a synergistic effect between *LRRK2* G2019S and ageing-dependent glial activation under a chronic subthreshold dose of LPS. The interaction between *LRRK2* G2019S, ageing, and low-grade systemic inflammation promoted PD-like features: (i) TH^+^ neuron loss in the SNpc and a marked reduction in DA striatal innervation; ii) motor deficits, and (iii) the up-regulation of several key proteins involved in mDAn dysfunction/death.

PD is a complex disorder that affects the central as well as the peripheral nervous system leading to a variety of motor and non-motor clinical symptoms. While the underlying causes remain unknown, converging evidence suggests a key role for gene/environment interactions in the initiation of the neurodegenerative events. Ageing and inflammation are among the key factors that may promote neuronal dysfunction and degeneration.^2,22,32,46,102^ LRRK2, the most frequent genetic cause of familial PD, is involved in the regulation of several ageing-related pathways including α-syn pathology, mitochondrial dysfunction as well as immune pathways and glia physiopathology.^24,38, 102–108^ Importantly, LRRK2 kinase activity is significantly increased in SN DA neuron and microglia in postmortem brain tissue from patients with idiopathic PD, indicating that LRRK2 can play a role in sporadic age-dependent PD.^11^ Several rodent models have been used to study LRRK2 biology and LRRK2 PD. LRRK2 transgenic mice generally exhibit mild phenotypes without a significant nigrostriatal degeneration nor motor deficits.^18–21, 109–111^ Overexpression (OE) of pathogenic *LRRK2* mutants, such as G2019S, R1441C, or R1441G, induces PD-like phenotypes, including age-dependent DA neuronal loss, disruption of dopamine homeostasis, L-DOPA responsive locomotor defects, and pathological accumulations of tau and α-syn.^15,112–117^ While OE models enable the investigation of PD neurodegenerative features, they are limited by potential overexpression-related artifacts. Previous studies have also examined the combined effect of ageing, environmental toxicity, and LRRK2-related genetic vulnerability. Chronic treatment of *LRRK2* R1441G knockin mice with the DA-specific environmental toxin rotenone causes increased locomotor deficits in mutant mice in the absence of significant nigrostriatal degeneration.^26^ These findings suggest that additional factors may interact with ageing and genetic vulnerability to trigger PD. One such factor could be inflammation.

To study the interaction between LRRK2 genetic vulnerability, ageing, and low-grade inflammation, we treated hemizygous *LRRK2* G2019S mice with low-dose systemic LPS. We selected two previously well-characterized temporal windows-young-adult and middle-aged.^67–69,72–75^ Middle age is recognized as a critical temporal window, whereby mDAns progressively become more vulnerable, and the microenvironmental SNpc milieu slowly shifts towards an “harmful” phenotype characterized by increased glial reactivity, oxidative stress, and the gradual loss of dopamine-mediated mitigation of inflammation.^22,46,102,118^ Based on our previously published findings and literature reports, coupled with pilot studies, we herein selected a very low, sub-threshold dose of 0.1 mg Kg^−1^, which induces a mild inflammatory response in young adult (3 M-old) as opposed to the exaggerated inflammatory response of aged (≥ 20 M-old) mice. ^66,68,76^ This dose did not induce sickness behavior (assessed by changes in body weight, food and fluid intake, and locomotor activity) when compared to NaCl injected mice. Such an LPS chronic regimen did not alter general health parameters studied nor life span.

Here, we show that chronic treatment of middle age *LRRK2* G2019S mice with low-dose LPS results in a significant recruitment of peripheral monocytes to the SNpc as well as CD4^+^ cells to the colon crypts. Interestingly, both age and chronic low-dose LPS synergistically induced LRRK2 expression in the SNpc and the spleen, a main reservoir of T lymphocytes, ^119^ with a more significant effect in G2019S mice. These data would be in line with a role of LRRK2 in sporadic PD.^11^ G2019S mice showed a significant infiltration of CD3^+^ cells within the SNpc as well as exacerbated astrocytosis and microgliosis both in the SNpc and the Str. Interestingly, key pro-oxidant and pro-inflammatory markers involved in SNpc DA degeneration were upregulated within the VM of G2019S mice, both at the mRNA and protein levels. This was accompanied by increased accumulation of α-syn, pS129α-syn, and pTau, which, in turn, could enhance immune responses and mDAn degeneration. Importantly, we show that peripheral immune activation precedes brain inflammation, macrophage/monocyte infiltration, and mDAn degeneration. Specifically, LPS treatment induced a proinflammatory phenotype in spleen macrophages and a significant dysregulation of peripheral cytokines, which was more prominent in G2019S mice. Furthermore, we show an enhanced CD4^+^ T-cell infiltration and α-syn aggregation in the colon of LPS-treated G2019S mice from 6 M on. These data suggest that the LRRK2 genotype is associated with an-age dependent gut immune cell infiltration and α-syn aggregation.

The present study does not establish a direct link between gut inflammation and proteinopathy with central inflammation and nigrostriatal degeneration. However, one hypothesis could be that *LRRK2* G2019S combined with a moderate chronic systemic inflammation triggers an age-dependent dysfunctional crosstalk of the gut-brain-axis, favoring the infiltration of pro-inflammatory CCR2^+^ peripheral monocytes overexpressing LRRK2 into the SNpc. Infiltrating cells and the peripheral immune cascade would interact and activate resident astrocytes and microglia. Studies are in progress to unveil whether and how gut α-syn pathology contributes to the progression of mDAn degeneration observed in this model.

The current findings suggest that genetic susceptibility promotes gut inflammation and α-syn aggregation that act as a driver of PD pathology. Several lines of evidence point to the impairment of the intestinal barrier function and colonic α-syn as a prodromal event in PD ^39–41,120^ i) the majority of patients experience non-motor symptoms involving the GI tract, such as chronic constipation and impairment of GI transit, several years before the onset of motor symptoms; ii) a- syn pathology is observed in the enteric nervous system of patients up to 20 years prior to diagnosis; and iii) increased intestinal permeability has been described in patients.^121^ Furthermore, changes in GI permeability and dysbiosis associated to a lowered serum LPS-binding protein in PD. ^122^ Interestingly, in the study of Kelly and coll. 2014, ^123^ a single, systemic high dose of 2.5 mg Kg^−1^ of LPS in young adult mice caused sequential increases in phosphorylated serine 129-α-syn and increased intestinal permeability in the colon starting 4 M after the acute LPS challenge, returning to baseline levels at 5 M.^123^ However, no nigrostriatal degeneration was observed. ^123^

Interestingly, we found a significant downregulation of the DA transcription factors *Nurr1* and *FOXA2* as well as a decrease of *TH* an *DAT* in G2019S VM under NaCl and/or LPS regimen, compared to their WT counterparts, suggesting increased vulnerability of G2019S mDAns. *Nurr1* and *FOXA2* are necessary for acquiring the mature DA phenotype and they also contribute to mDAn maintenance, protection, α-syn expression, and immunomodulation. ^46,102,124–130^ The significant reduction of *Drd2* levels in the VM of ageing G2019S mice under LPS, as compared to their WT counterparts, highlights the critical role of dopamine signaling in counteracting oxidative stress and inflammation in the aged parkinsonian brain.^102^ Notably, D2-like receptor subtypes, coupled to Gαi/o, suppress cAMP activity with an inhibitory effect upon DA binding. ^131^ Elevated cAMP activates PKA and phosphorylates cAMP-response element binding protein (CREB), which disrupts NF-κB homeostasis and dysregulates the inhibition of the inflammatory response.^132^ DrD2 are expressed in astrocytes, microglia, peripheral immune cells as well as intestines.^102, 132–134^ Hence, the decrease of DrD2 could contribute to the inflammation-dependent nigrostriatal neurodegeneration observed in LRRK2 G2019S mice. Interestingly, recent work shows that intestinal DrD2 are necessary for neuroprotection against the PD neurotoxin 1-Methyl-4-phenyl-1,2,3,6-tetrahydropyridine (MPTP)-induced mDAns death^135^. The selective ablation of Drd2, but not Drd4, in the intestinal epithelium, causes a more severe loss of mDAns following MPTP challenge that is associated with a dysfunction of the gut microbiota.^135^ Future studies will address whether intestinal DrD2 contribute to the inflammatory intestinal milieu in the G2019S-ageing-inflammatory model.

Another possible mechanism involved in mDAn loss in G2016S mice could be the increase of LRRK2 protein, which in turn regulates tau phosphorylation through activation of GSK-3β^84–86^. Active pTyr^216^-GSK-3β signalling plays a key role in mDAn death and it may be involved in region-specific phosphorylation and accumulation of tau and α-syn.^72, 87–91, 136^ Furthermore, dysfunctional GSK-3β signalling modulates the neuron-glia crosstalk. ^46,83,89,102,137^ A critical role of active GSK-3β may also be relevant for LRRK2 PD. ^84,85,138–140^

With regards to the astrocytic response, middle-age was confirmed as the most vulnerable age-group in both genotypes, with LRRK2 G2019S mice showing an enhanced astrocytic response. While astrocytes play key neuroprotective roles,^46,75,89,102,103,141,142^ with age, they become dysfunctional and they can contribute to the increased mDAn vulnerability.^46,89,102,103,141,142^ Hyper-active astrocytes and increased expression of inflammatory cytokines and other toxic mediators are prominent features of PD.^33,46,75,89,102,103,141,142^ Several abnormalities of astrocyte function were recently reported in G2019S astrocytes, including defects in the clearance of α-syn.^103,143–145^ However, the contribution of astrocytes to LRRK2 PD pathophysiology *in vivo* is still unknown. In our study, LPS-treated WT mice displayed healthier TH neuronal cell bodies and longer TH^+^ neurites with close interactions with GFAP^+^ astrocytic processes. On the contrary, G2019S mice showed TH^+^ neuron atrophy associated to poor contact with GFAP^+^ astrocytes. These data would suggest that WT astrocytes might contribute to support the vulnerable mDAn upon a subthreshold inflammatory trigger, while G2019S astrocytes likely shift towards a reactive harmful phenotype. ^46,75,102,141,142,146,147^ In experimental models of multiple sclerosis, inflammatory cytokines mediate local astrocyte-dependent mechanisms of immune cell trafficking within the CNS. ^148^ Recent work shows that the degree of inflammatory activity in MS patients at disease onset induces a reactive state in astrocytes that triggers neuronal damage.^149^ Our data suggest that local astrocyte-macrophage/monocyte interactions in LRRK2 G2019S mice promote astrocyte-dependent mechanisms of immune cell trafficking within the CNS during ageing and low-grade inflammation. However, further studies carried out in PD-specific brain regions, peripheral tissues and blood are clearly needed to explore in this LRRK2-inflammatory model, “in situ” cell-cell interactions and novel signaling mechanisms that might be useful to develop ageing-specific therapeutic targets^150^.

With regard to microglia, both *in vitro* and *in vivo* studies show that *LRRK2* modulate microglia inflammatory responses.^53,55,56,151–153^ Recent studies suggest that LRRK2 levels in microglia may not have a direct effect on neuroinflammation in PD, as *in vivo* injection of a high acute dose of LPS failed to increase microglia LRRK2 protein levels in R1441C and G2019S young adult mice^70^. Similar results have been observed *ex vivo*, with neither LPS ^154^ or priming with a-syn pre-formed fibrils ^155^ increasing LRRK2 protein expression in cultured murine microglia. Recent findings from Langston and co, implicated a genetically-mediated cell-type and region-specific effect of LRRK2 in microglia ^156^. Our results strongly indicate that both ageing and a sub-threshold chronic inflammation in conjunction with LRRK2 genetic vulnerability promote peripheral and central immune dysregulation.

In rodent models of PD, peripheral immune cells interact with resident immune cells in diseased brain to promote TH neuron loss.^42,44,45,48,157^ For example, peripheral monocyte entry is required for α-syn induced inflammation and neurodegeneration,^100^ and reciprocally, pathological α-syn was shown to recruit LRRK2 expressing pro-inflammatory monocytes to the brain.^158^ LRRK2 is highly expressed in circulating and tissue immune cells and it is upregulated following recognition of microbial structures.^51^ However, it has been reported that acute endotoxemia in 3 M-old LRRK2 mutant and WT mice does not trigger T-cell infiltration in the brain.^70^ In contrast, we show that a low-dose chronic inflammation in ageing LRRK2 mice triggers a massive monocyte/macrophage infiltration associated to a robust microglia activation. Both gene expression and protein analyses revealed the up-regulation levels of inflammatory and oxidative stress markers within the VM of LRRK2 G2019S mice. This was paralleled by an increase of α-syn phosphorylated at Ser-129, which is the dominant pathological modification of α-syn both in familial and sporadic PD. ^91,92^ Brain-engrafting pro-inflammatory monocytes have been implicated in DA neurodegeneration in PD models. Remarkably, in 6M-old mice, we did not find CD3^+^ T cell brain infiltration in none of the genotypes under low-grade inflammation, whereas CD3^+^ cells were clearly stained in spleen sections of the same age and treatment group. Starting at 10 M on, we observed a significant increase in CD3^+^ /LRRK2^+^ cells in G2019S SNpc compared to WT counterparts. Moreover, LRRK2 G2019S synergized with LPS and IFN-γ in SPMs to promote greater levels of TNF-α and IL-6 release starting at 6 M. Given the recently recognized role of the brain-spleen axis,^159,160^ these data would suggest a potential early role of the peripheral immune system in LRRK2-related neurodegeneration. In line with these data, we found a significant increase in the serum levels of IL-6, IFN-γ, IL1-beta, IL-2, IL-4 and IL-17 in G2019S LPS-treated mice at 6 M. These results indicate an overactivation of the peripheral immune system in G2019S mice subjected to low-grade inflammation during ageing. Peripheral cytokines could activate brain endothelial cells and increase the blood-brain barrier permeability, which in turn facilitates immune cell infiltration into the brain parenchyma. ^161^ Furthermore, given that TLR4 is present in the circumventricular organs that lack the blood brain barrier,^162–165^ in perivascular macrophages, microglia, astrocytes, and in trigeminal and nodose ganglia, ^166,167^ these processes could amplify peripheral LPS effects in the brain.

## 5 CONCLUSION

We established a novel translational model of LRRK2 PD, showing an interaction between the most crucial risk factors involved in both idiopathic and familial PD. We show that low-dose inflammation during middle-age triggers mDAn degeneration in LRRK2 G2019S mice, accompanied by astro- and microgliosis as well as macrophage/monocyte CNS infiltration. Brain inflammation and degeneration are preceded by the dysregulation of peripheral cytokines as well as increased CD4^+^ T-cell infiltration and α-syn aggregation in the colon. LRRK2 G2019S exacerbates the recruitment of peripheral monocytes and CD3^+^ T cells to the brain in response to low-grade inflammation and ageing, likely driving astrocyte and microglial activation and accumulation of pathological forms of α-syn (**Figure 10**). Importantly, these results suggest an early role of the peripheral immune system, including the gut, in PD pathogenesis. This novel preclinical model of LRRK2 PD will enable the identification of key players involved in mDAn degeneration, both at the central and peripheral level, and it will be useful to explore age- and immune-related biomarkers and targets with therapeutical relevance for both sporadic and LRRK2 PD.

**Figure 10.**
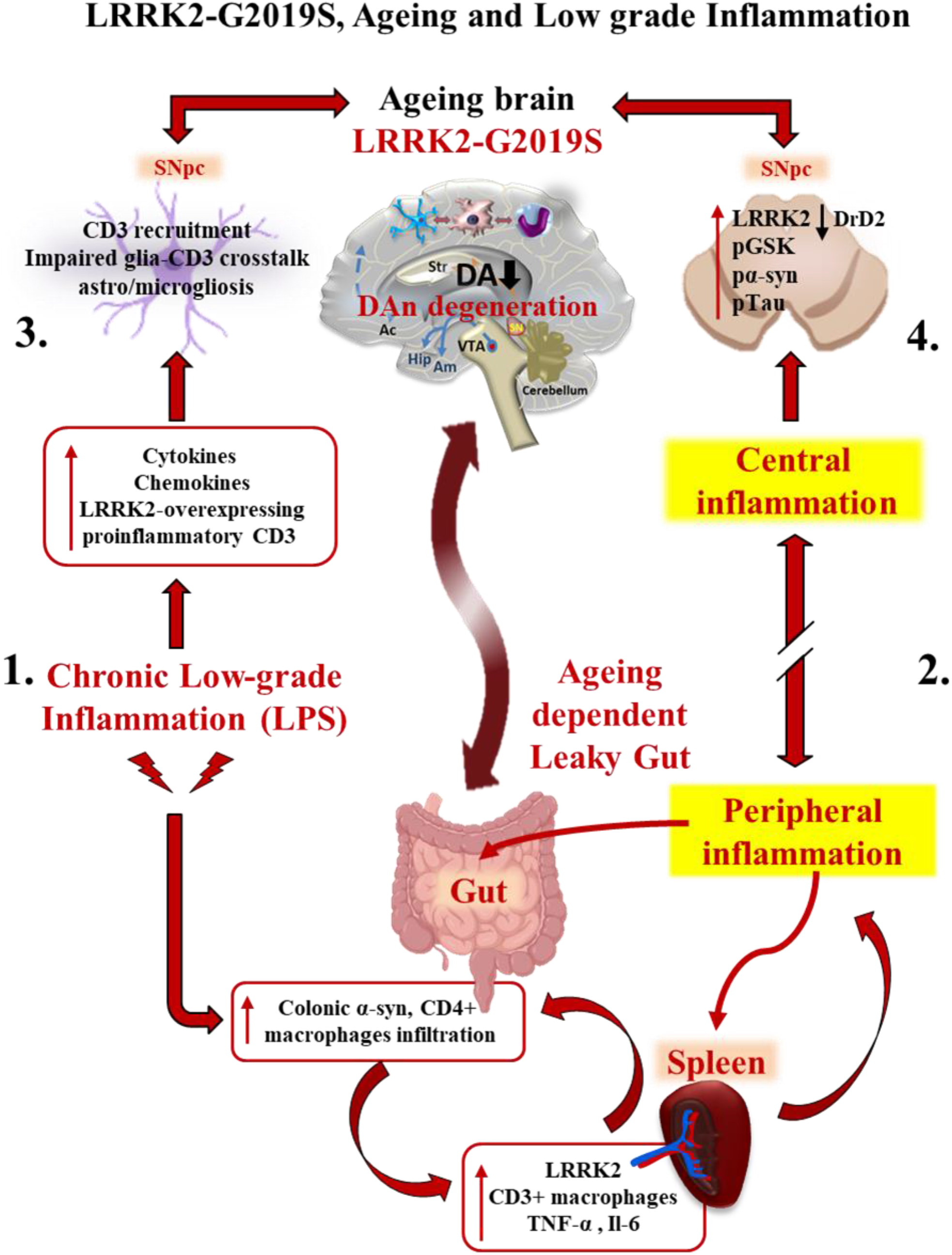
LRRK2 G2019S interacts with low-grade inflammation and ageing to promote peripheral inflammation and dopaminergic neuron degeneration. Schematic drawing showing the mechanisms involved in LRRK2 G2019S-mediated midbrain neuron (mDAn) degeneration in the ageing brain. **1**. LRRK2 G2019S interacts with low-grade inflammation (subthreshold dose of LPS) promoting early CD4+ T cell infiltration and α-syn aggregation in the gut as well as **2**. dysregulation of the peripheral immune system (changes in peripheral cytokines and activation of spleen macrophages). **3**. Peripheral immune changes may promote macrophage/monocyte infiltration in the CNS, astro- /microgliosis, and CD3+ cell infiltration; **4**. LRRK2 G2019S synergizes with ageing and low-grade chronic inflammation to promote the up-regulation of several key factors involved in mDAn dysfunction/death.

## Acknowledgements

This project was supported by the “Network of Centres of Excellence in Neurodegeneration (CoEN) 2017, Pathfinder III project”, “Investigating the interaction between ageing and immune dysfunction in LRRK2 Parkinson’s disease” to MD, (University of Tübingen-Germany), BM (OASI-IRCCS-Troina, Italy). CG had a Coen Grant fellowship (OASI-IRCCS-Troina). We are grateful to CAPiR (Center for Advanced Preclinical *in vivo* Research) Team, of the University of Catania, for maintenance and care of animals. The authors acknowledge the laboratory of Neuropharmacology (BM, OASI-IRCCS-Troina), the Pharmacology Laboratory section (BM, BIOMETEC) and the technical assistance at Bio-nanotech Research and Innovation Tower (BRIT, BIOMETEC). The research program also received support from the Italian Ministry of Health (Cur. Res. projects 2018-2022, to B.M.), from University of Catania (PIACERI and PhD program in Biotechnology, to CG and BM), the Helmholtz Association Young Investigator Award (VH-NG-1123, to MD).

## 11 ETHICS APPROVAL AND CONSENT TO PARTICIPATE

All animal experiments were performed in strict accordance with the Guidelines for the Care and Use of Laboratory Animals (NIH Publication, revised 2011), which were also approved by the Animal Care and Use Committee of OPBA of the University of Catania.

## 12 AUTHOR CONTRIBUTIONS

All authors have contributed to the conception and design. All authors have contributed to Material preparation, data collection and analyses were performed by Bianca Marchetti, Michela Deleidi, Carmela Giachino, and Cataldo Tirolo. The first draft of the Manuscript was written by Bianca Marchetti and Michela Deleidi and all authors commented on previous versions of the manuscript. All authors read and approved the final manuscript.

### DATA AVAILABILITY STATEMENT

All data that support the findings in this study are available from the corresponding author upon reasonable request.

---

**Supplemental Figure 1.**
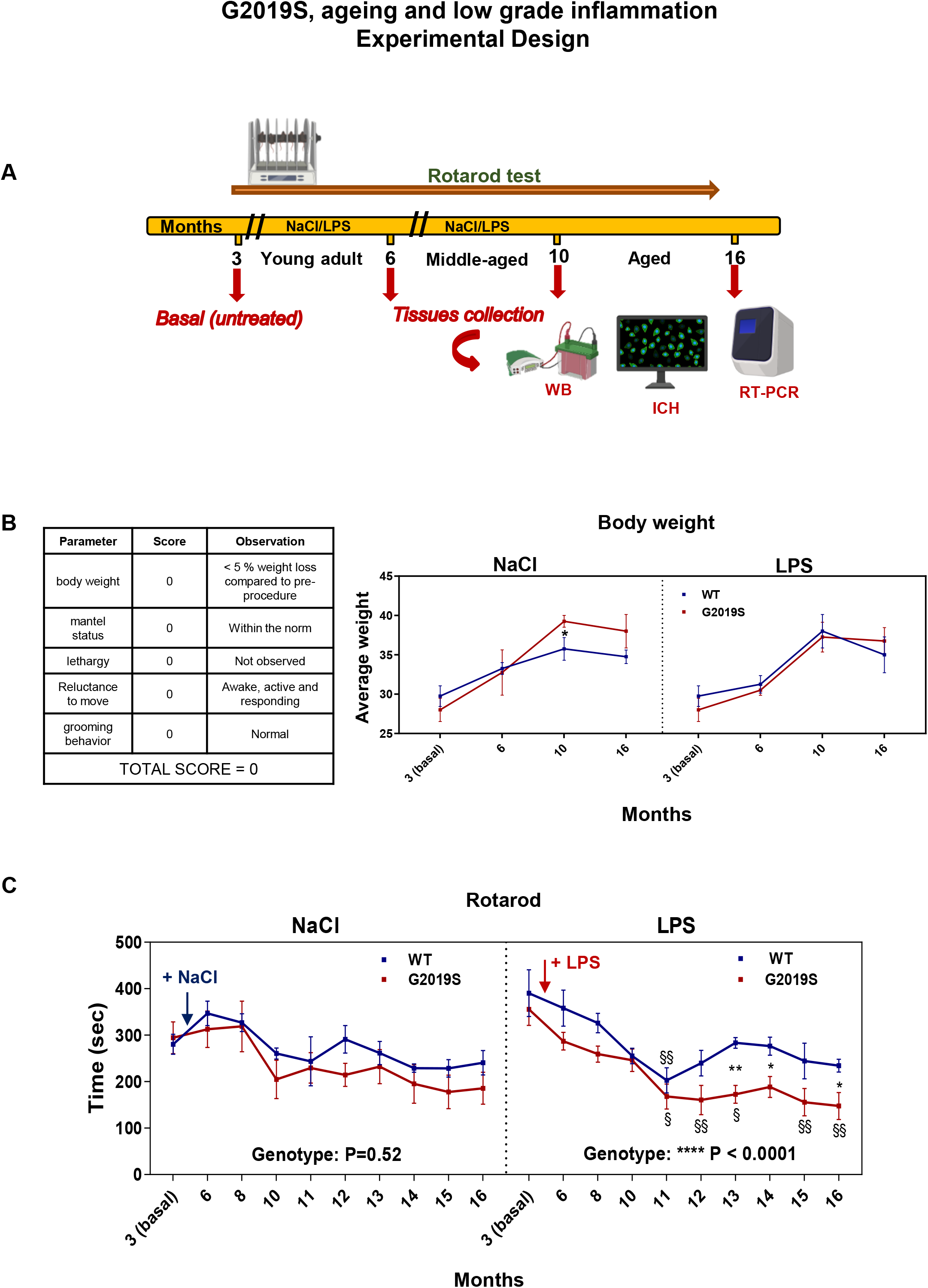

**Supplemental Figure 2.**
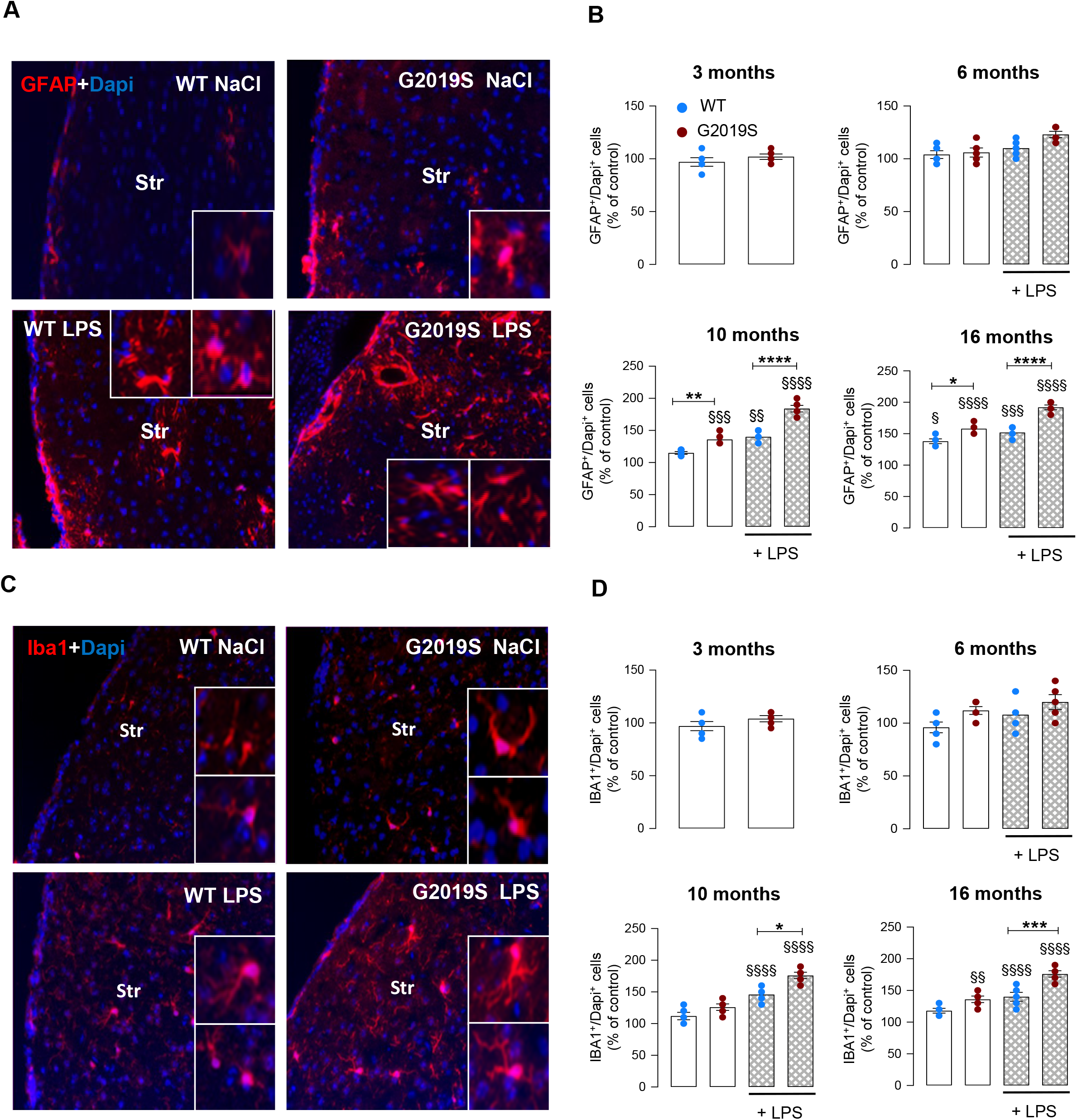

**Supplemental Figure 3.**
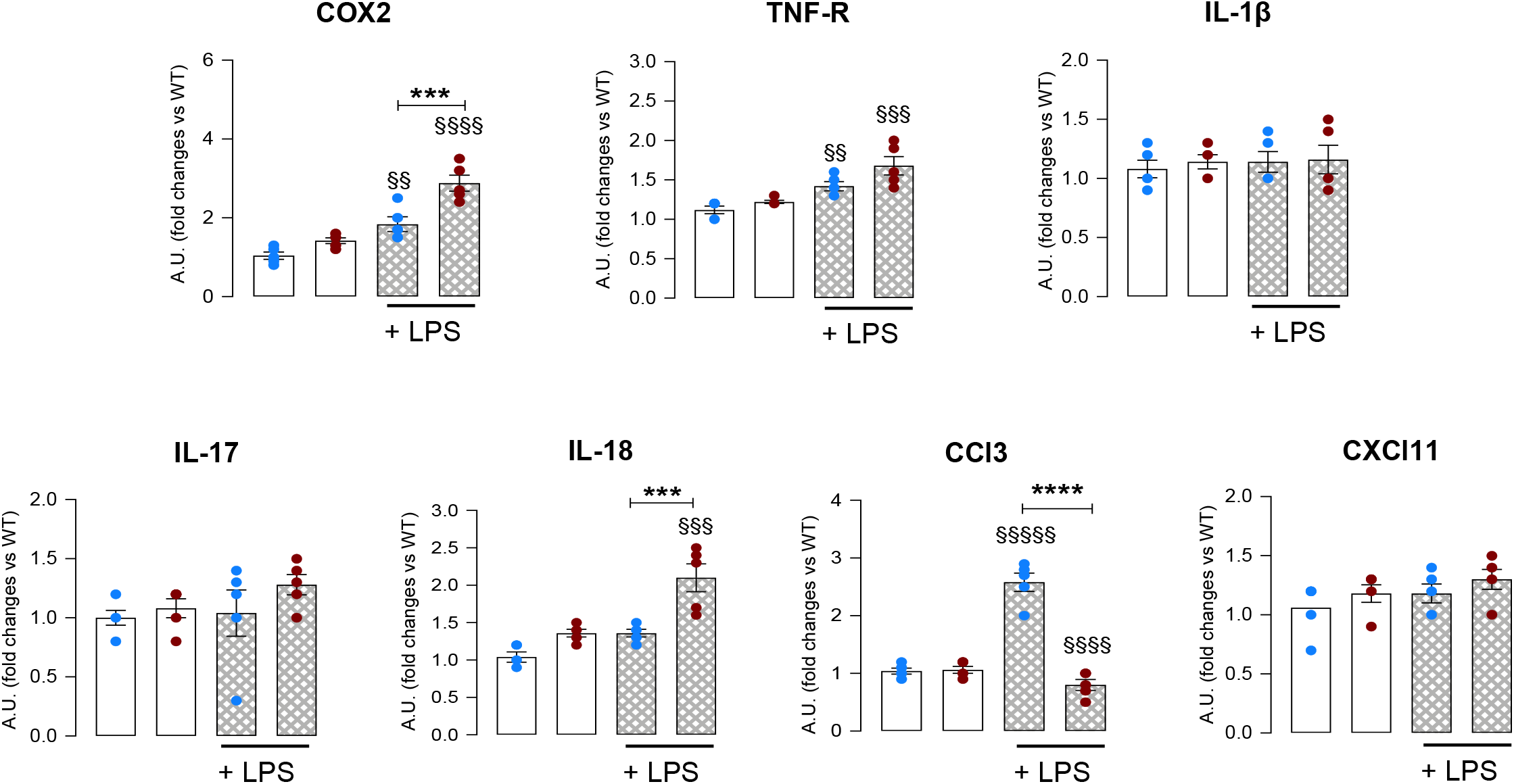

**Supplemental Figure 4.**
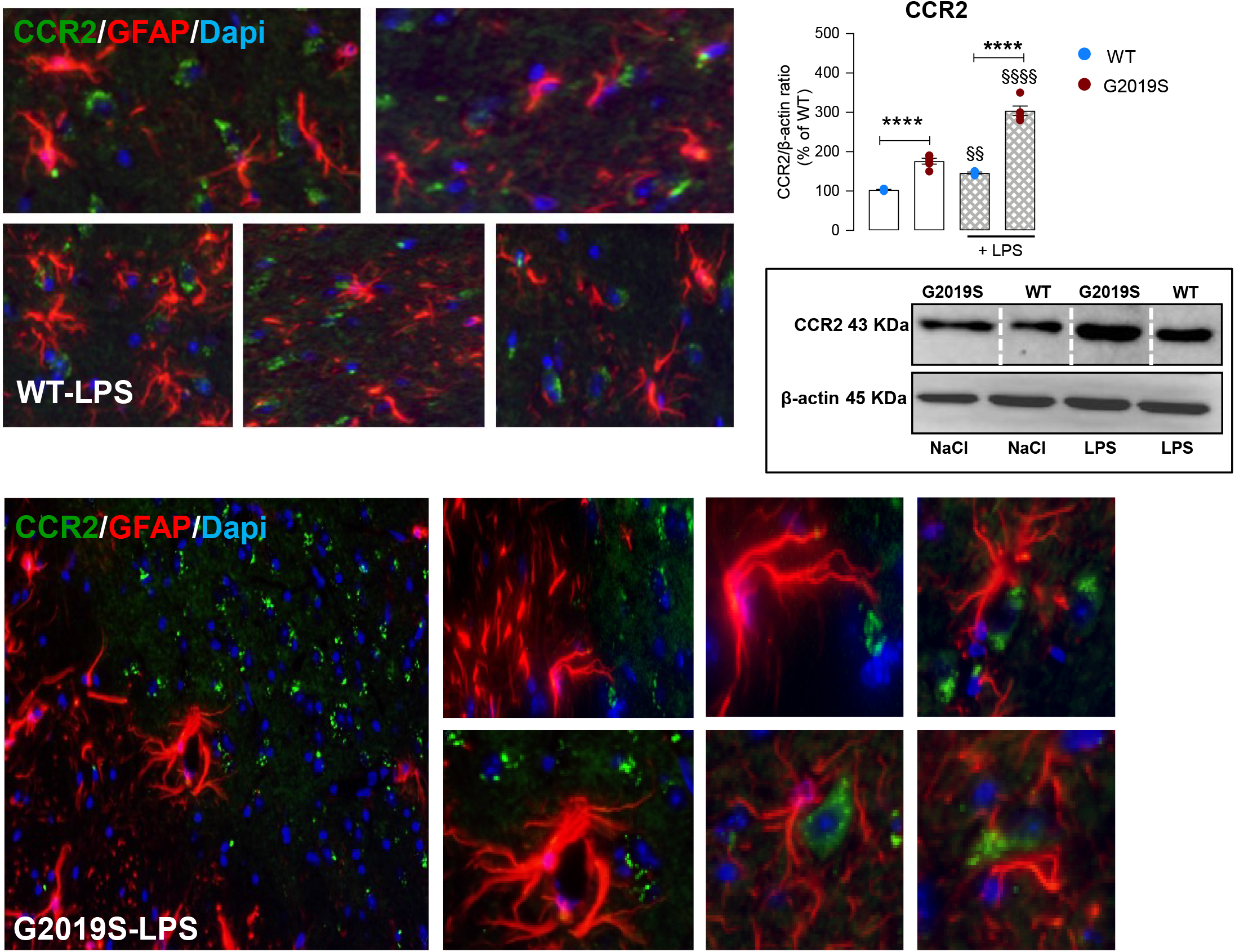

## SUPPLEMENTARY INFORMATIONS

### METHODS

#### IMMUNOHISTOCHEMISTRY

The mice (5-6 mice/genotype/age-group/treatment) were anesthetized and transcardially perfused with 0.9% NaCl, followed by 4 % paraformaldehyde in phosphate buffer (pH 7.2 at 4°C), the brains and peripheral tissues carefully removed and post-fixed for 2-4 hrs, in 4% paraformaldehyde in phosphate buffer (pH 7.2) and placed in 15% sucrose in the solution of phosphate buffer overnight at 4°C. Tissues were frozen and stored at −80° C until further analyses. For the brain, serial coronal sections (14 μm-thick), encompassing the striatum (Bregma 1.54 to bregma −0.46) and the SNpc (Bregma −2.92 to bregma −3.8 mm) according to Franklin and Paxinos (1997) were collected, mounted on poly-L-lysine-coated slides and processed as previously described in full details (L’Episcopo et al. 2011b). The pre-absorbed primary antibodies used are shown in *Supplemental Table 1*. Sections were washed extensively and incubated with fluorochrome (FITC, CY3, CY5)-conjugated species-specific secondary antibodies for immunofluorescent detection. TH immunoreactivity was also detected using biotinylated secondary antibodies (Vector Laboratories) and diaminobenzidine (DAB, Vector Laboratories) as the developing agent. Cresyl violet was used to visualize Nissl substance. In all of these protocols, blanks were processed as for experimental samples except that the primary antibodies were replaced with PBS. After overnight incubation, sections were rinsed and incubated in darkness for 2 h with CY3-conjugated donkey anti-rat, donkey anti-rabbit, and donkey anti-goat antibodies (1:200; Jackson ImmunoResearch), mounted on glass slides and coverslipped with glycerol-based mounting medium.

#### CELL COUNTS, QUANTIFICATION OF IMMUNOHISTOCHEMISTRY AND IMAGE ANALYSIS

Dopaminergic neuronal counts were determined by serial section analysis of the total number of TH^+^ neurons in the right and left SNpc throught the entire extent of the SNpc using DAPI or PI as nuclear markers, as previously described (L’Episcopo et al 2011b). Briefly, at least three sections were obtained from each animal representing each of the five representative planes from −2.92 to −3.8 mm relative to bregma according to the stereotaxic coordinates (Franklin and Paxinos (1997). Total numbers of TH- and cresyl violet (CV)-stained neurons in adjacent tissue sections were estimated in parallel to validate TH^+^ neuron survival (Baquet et al. 2009; L’Episcopo et al. 2018). A total of ≥ 5 mice/group/time-point was analyzed in a blind fashion. Each midbrain section was viewed at low power (X 10 objective) and the SNpc was outlined and delineated from the ventral tegmental area immunoreactive neurons by using the third nerve and cerebral peduncle as landmarks. For fluorescence microscopy, a confocal laser scanning microscope LEICA TCS-NT (Version 2.5, Build 1227, Leica Microsystems GmBH, Heidelberg, Germany, equiped with image analysis software), with an argon/krypton laser using 10 X, 20 X, and 40 X and 100 X (oil) immersion objectives, was used. TH immunoreactivity was also detected using biotinylated secondary antibodies and CV was used to visualize Nissl substance. Estimates of total TH^+^ and CV-stained neurons in the SNpc were calculated using Abercrombie correction (Baquet et al., 2009; L’Episcopo et al. 2011b; 2018). Briefly, the Abercrombie correction considers the number of sections collected, the interval between sections, and the thickness of each section to estimate the total number of neurons (cresyl violet) or TH+ cells within the entire SNpc. The Abercrombie values are calculated by multiplying the total neuronal count for each of these factors. The total number of TH^+^Nissl^+^ neurons was estimated at the different time-intervals studied. Results of the right and left SN were thus added to generate a total TH^+^ SNpc neuron count. Treatment groups were averaged and represent the means ± SEM. Differences were analyzed by ANOVA followed by Tukey’s multiple comparison or Bonferroni-post hoc tests, and considered significant when *P* < 0.05.

Striatal TH- and dopamine transporter (DAT)-immunoreactive (IR) fiber staining was assessed in n = 3 coronal sections at three levels (bregma coordinates: + 0.5, + 0.86, and 1.1 mm, respectively) of caudate-putamen (CPu), in ≥ 5 mice/group/time. In all cases of immunohistochemical quantification, analyses were performed by an individual unaware of the experimental treatments. Fluorescence intensity (FI) of TH-staining above a fixed threshold using the corpus callosum for background subtraction (Burke et al., 1990; L’Episcopo et al. 2011b; 2018). Measurements of FI were carried out by computer-assisted image analysis software (LEICA), and changes in average FI (mean ± SEM) expressed as percentage (%) of saline-injected controls. Cell counts were obtained for ameboid IBA1/Dapi^+^ or Mac-1^+^/Dapi^+^ reactive microglial (Gennuso et al. 2004; L’Episcopo et al. 2011b;c) cells and GFAP^+^ Dapi^+^ astrocytes, averaged for each animal and the mean number of cells per mm^2^ per animal was estimated. A comparable countable area ranging from 1.90 mm^2^ to 2.00 mm^3^ was analyzed in the different groups. Results are expressed as % of saline-injected controls.

#### TISSUE PREPARATION FOR NEUROCHEMICAL/MOLECULAR ANALYSIS

At the indicated time-intervals, mice were sacrificed by cervical dislocation, the brains rapidly removed and immediately placed on ice-cold saline. The right and left striata and ventral midbrain (which included the SNpc) were then dissected on an ice-cold plastic dish and processed as described.

#### GENE EXPRESSION ANALYSIS

Gene expression was performed on tissue samples of the ventral midbrain (containing the substantia nigra pars compacta, SNpc) (L’Episcopo et al. 2011b; 2018).

##### RNA Extraction

At due time points, tissue samples were homogenized in 1ml of QIAzol Lysis Reagent (Qiagen, #79306) using a rotor-stator homogenizer. Total RNA was isolated from homogenized tissue samples using RNeasy Lipid Tissue Kit (Qiagen, #74804) including Dnase digestion. At the end, RNA samples were redissolved in 30 μl of RNase-free water and their concentrations were determinated spectrophotometrically by A_260_ (Nanodrop-ND 1000).

##### Reverse Transcription

cDNA synthesis were performed using Retroscript Kit (Ambion #1710, Austin, Texas) according the manufacturer’s instructions. 50 μl of water solution containing 0.5 μg of each pool were added to an equal volume of 2X TaqMan Universal PCR Mastrer Mix (Applied Biosystems).

##### Real Time PCR

Real-time quantitative PCR was performed using Taqman™Assay Reagents on an Step One Detection System (Applied Biosystems) according to manufactures protocol. Tissue samples were processed as above. Cell samples were lysed in lysis buffer (Qiagen) and stored at −80°C until the RNA was extracted following manufacture’s instructions. Residual genomic DNA was removed by incubating with DNase I, RNase-free (Qiagen) and eluted from the RNeasy mini columns with RNase-free water. The amount of total RNA was quantified using a NanoDrop ND-100 (Nano Drop Technologies) and the cDNA was synthesized from 2 μg of total RNA using the Retroscript Kit (Ambion). After purification using QIAquick PCR Purification kit (Qiagen), 250 ng of cDNA were used for Real-time PCR using pre-developed Taqman Assay Reagents (Applied Biosystems). Real-time quantitative PCR was performed with Step One Detection System (Applied Biosystems) according to manufacturer protocol. The assay IDs are reported in *Supplemental Table 2*. We used the housekeeping gene, β-actin, as normalizer and embrionic mouse brain as calibrator. Quantification of the abundance of target gene expression was determined relative to β-actin with respect to the control group by using the delta delta C_t_ (2^−ΔΔCt^) comparative method, the results expressed as arbitrary units (AU). Relative fold changes over WT in the respective treatment groups are indicated.

#### WESTERN BLOT ANALYSIS

Protein extracts were prepared as described in full details (L’Episcopo et al. 2016; 2018). The antibodies were used in this study are presented in *Supplementary Table 1*. The tissue samples were homogenized in lysis buffer (0.33 M sucrose/8 mM Hepes, pH 7.4 and protease inhibitors) and quantified using the BCA protein determination method (Bio-Rad, Hercules, CA). Protein samples were diluted to equivalent volumes containing 20 μg of protein and boiled in an equal volume of Laemli SDS boiling buffer (Sigma) for 10 min. Samples were loaded into a 9-12% SDS-polyacrilamide gel and separated by electrophoresis for 3 h at 100 V. Proteins were transferred to polyvinylidene difluoride membrane (Amersham Biosciences, Piscataway, NJ) for 1.5 hr at 300 mA. After blocking of nonspecific binding with 5% non-fat dry milk in TBST, the membranes were then probed with the indicated primary antibodies (*Supplementary Table 1*). After incubation at room temperature for 1 hr, membranes were washed and treated with appropriate secondary antibodies conjugated with horseradish peroxidase (HRP) and blot were exposed onto radiographic film (Hyperfilm; Amersham Bioscience). Membranes were reprobed for β-actin immunolabeling as an internal control. The bands from the Western blots were densitometrically quantified on X-ray films using a software to determine the levels of immunoreactivity (ImageQuantity One). The data from experimental band were normalized to β-actin. Values of phosphorylated GSK-3β (pTyr^216^ GSK-3β); phosphorylated α-syn (pSer^129^ α-syn) and phosphorylated tau (pSer^396^ tau) were normalized for each respective control (total GSK-3 β, α-syn, and tau, respectively), before statistical analysis of variance and values expressed as percent changes (%) of WT controls (L’Episcopo et al. 2016). Dashed lines (in white) indicate discontinuous bands (nonsequential lanes) taken from the same blot, at the same molecular weight (mass - kDa) in order to better represent the mean signal from all values for that particular group, and should not be confused with continuous bands (side by side) from same blot. Corresponding control bands (loading controls) match experimental bands.

#### *EX VIVO* CULTURES OF SPLEEN MACROPHAGES AND BV2 CELLS

Spleens were dissected from abdominal cavity and filtered through a 40-μm nylon strainer. Red cell lysis buffer was used to remove red cells. A single splenic cell suspension then was obtained (Marchetti et al. 2002). Cells were cultured in Roswell Park Memorial Institute (RPMI) medium 1640 RPMI 1640 (BioConcept 1-41F01-I) supplemented with 10% FBS, 2mM L-Glutamine and antimicrobials (Penicillin-Streptomycin Pen 10’000 IU/ml Strep 10 mg/ml and amphotericin B (250 μg/ml BioConcept). Mouse microglia BV2 cells (Henn et al. 2009) from Elabscience (No.: EP-CL-0493) were cultured in parallel for each spleen culture preparation and served as controls. Briefly, BV2 cells were maintained in Roswell Park Memorial Institute (RPMI) medium 1640 supplemented (BioConcept 1-41F01-I) with 10% FBS (FBS-02-0500), 2mM L-Glutamine 5-10K50-H) and antimicrobials (Penicillin-Streptomycin Pen 10’000 IU/ml Strep 10 mg/ml and amphotericin B (250 μg/ml) (BioConcept 4-01F00-H). Spleen macrophages (SPMs) differentiate into the M1 phenotype after stimulation with LPS (100 ng/ml) ± IFN-γ (10 ng/ml) (Orihuela et al. 2016). For BV2 stimulation, RPMI was replaced by Dulbecco’s Modified Eagle Medium (DMEM) High Glucose. (BioConcept, 1-26F03-I).

#### TNF-α AND IL-6 DETERMINATION BY ELISA

To monitor M1 status, of SPMs and BV2 cells, we evaluated Th1 and pro-inflammatory cytokines, TNF-α (MTA00B) and IL-6 (M6000B) determined using enzyme-linked immunosorbent assay (ELISA) kits (ELISA Development System; R&D Systems, McKinley Place, MN, USA) following the manufacturer’s protocol. This assay employs the quantitative sandwich enzyme immunoassay technique (L’Episcopo et al. 2018).

---

**Table 1.** List of antibodies used for immunohistochemistry and western blot

| **Antibody name** | **Company** | **Dilution** |
| --- | --- | --- |
| CCR2, goat polyclonal | Thermo Fisher Scientific | 1:500 |
| CD11b, rabbit | Abcam | 1:1000 |
| CD11b, rat | Biologend | 1:500 |
| CD3, mouse | Santa Cruz Biotechnology | 1:500 |
| CD4, rat | Bio-Rad | 1:300 |
| CD68, rat | Bio-Rad | 1:300 |
| DAT, rabbit | Chemicon | 1:200 |
| GFAP, mouse | Sigma | 1:250 |
| GFAP, rabbit | Chemicon | 1:1000 |
| GSK-3β, rabbit | Santa Cruz B. | 1:200 |
| HuD+HuC, rabbit | Abcam | 1:400 |
| Iba-1, goat polyclonal | Abcam | 1: 200 |
| Iba-1, rabbit polyclonal | Wako | 1:500 |
| iNOS, mouse | Santa Cruz | 1:200 |
| LRRK2, rabbit | clone 41-2, Abcam by MJFF | 1:1000 |
| MAP2, rabbit | Cell Signaling | 1:500 |
| NeuN, rabbit | Abcam | 1:1000 |
| Nurr1, goat polyclonal | R&D | 1:300 |
| phospho-alpha-Synuclein (Ser<sup>129</sup>),<br>mouse monoclonal (clone LS4-1B1) | MilliporeSigma | 1:1000 |
| phospho-GSK-3β (p-Tyr<sup>216</sup>), rabbit | Abcam | 1:500 |
| Phospho-LRRK2-S<sup>935</sup>, rabbit | Abcam by MJFF | 1:1000 |
| phospho-Tau (pSer<sup>396</sup>), rabbit<br>polyclonal | Santa Cruz Biotechnology | 1:200 |
| phospho-Tau (pThr<sup>181</sup>) | Santa Cruz Biotechnology | 1:200 |
| Tau, mouse | Invitrogen | 1:200 |
| TH, mouse | Boehringer Mannheim | 1:200 |
| TH, rabbit | Pel-Freez | 1:200 |
| TH, rabbit | Chemicon | 1:200 |
| TH, sheep | Pel-Freez Biologicals | 1:200 |
| TH, sheep | Pel-Freez Biologicals | 1:1000 |
| TUBB3, mouse | Biologend | 1:400 |
| α-syn (C-20)-R, rabbit | Santa Cruz Biotechnology | 1:200 |
| α-syn aggregate, MJFR-14-6-4-2,<br>rabbit | Abcam | 1/5000 |
| β-actin, rabbit | Cell Signaling | 1:1000 |
| β-actin, rabbit or mouse | Abcam | 1:2000 |

**Table 2.** List of probes and IDs used for quantitative Real time PCR

| **PROBES** | **IDs** |
| --- | --- |
| Casp6 | Mm00438053-m1 |
| CCl3 | Mm 00441258-m1 |
| CxCl10 | Mm 00445235-m1 |
| CxCl11 | Mm00444662-m1 |
| Drd1a | Mm01353211-m1 |
| Drd2 | Mm00438545-m1 |
| Foxa2 | Mm01976556-s1 |
| GAPDH | Mm99999915-g1 |
| Gfap | Mm 00546086-m1 |
| Gp91phox | Mm01287743-m1 |
| GSK3b | Mm00444911-m1 |
| IL-10 | Mm12088386-m1 |
| IL-17 | Mm00439618-m1 |
| IL-18 | Mm00434226-m1 |
| Il1b | Mm00434228-m1 |
| IL-4 | Mm00445259-m1 |
| Itgam (Mac1) | Mm 00434455-m1 |
| LRRK2 | Mm00481934_m1 |
| Map2 | Mm00485231-m1 |
| Nfkb1 | Mm00476361-m1 |
| Nos 2 | Mm 00440485-m1 |
| Nurr 1 | Mm 00443056-m1 |
| Slc6a3 (Dat) | Mm 00438388-m1 |
| Tau (Mapt) | Mm00521988-m1 |
| Th | Mm 00447546-m1 |
| Tnf | Mm00443258-m1 |
| Tnfrsf1b | Mm00441875-m1 |
| α-syn (SNCA) | Mm00458965-m1 |

